# mixMC: a multivariate statistical framework to gain insight into Microbial Communities

**DOI:** 10.1101/044206

**Authors:** Kim-Anh Lê Cao, Mary-Ellen Costello, Vanessa Anne Lakis, François Bartolo, Xin-Yi Chua, Rémi Brazeilles, Pascale Rondeau

**Affiliations:** The University of Queensland Diamantina Institute, Translational Research Institute, Brisbane, QLD 4102, Australia; Institut de Mathématiques de Toulouse, UMR CNRS 5219 INSA Université de Toulouse, Toulouse, France; Queensland Facility for Advanced Bioinformatics, The Institute for Molecular Bioscience, Brisbane, QLD 4072, Australia; Danone Nutricia Research, Palaiseau Cedex, 91767, France

## Abstract

Culture independent techniques, such as shotgun metagenomics and 16S rRNA amplicon sequencing have dramatically changed the way we can examine microbial communities. Recently, changes in microbial community structure and dynamics have been associated with a growing list of human diseases. The identification and comparison of bacteria driving those changes requires the development of sound statistical tools, especially if microbial biomarkers are to be used in a clinical setting.

We present **`mixMC`**, a novel multivariate data analysis framework for metagenomic biomarker discovery. **`mixMC`** accounts for the compositional nature of 16S data and enables detection of subtle differences when high inter-subject variability is present due to microbial sampling performed repeatedly on the same subjects but in multiple habitats. Through data dimension reduction the multivariate methods provide insightful graphical visualisations to characterise each type of environment in a detailed manner.

We applied **`mixMC`** to 16S microbiome studies focusing on multiple body sites in healthy individuals, compared our results with existing statistical tools and illustrated added value of using multivariate methodologies to fully characterise and compare microbial communities.

## I. Introduction

The human gut microbiome contains a dynamic and vast array of microbes that are essential to health and provide important metabolic capabilities. Until recently, studying these complex communities has been difficult and generally limited to classical phenotypic techniques Clarridge (2004); Huse et al. (2010). With the improvement of high-throughput sequencing technology, the ability to profile complex microbial communities without the need to individually culture organisms has increased dramatically. These sequencing methods range from RNA sequencing (RNA-seq), chromatin immunoprecipitation sequencing (ChIP-seq), metagenomic and 16S rRNA gene amplification analysis of microbial populations. 16S rRNA sequencing in particular has substantially changed our understanding of phylogeny and microbial diversity. The analysis of this commonly used ribosomal RNA gene is quickly becoming a staple for profiling of microbial communities from soil to humans. With this sequencing technique, hypervariable regions within the gene are amplified, sequenced, and clustered into operational taxonomic units (OTU). Taxonomic classification of representative sequences from each cluster is then aligned against a database of previously characterised 16S ribosomal DNA (rDNA) reference sequences to indicate the ‘species’ or unit of interest.

With the rapid development of sequencing technologies, drop in price and increase in sample size, more than ever is being asked of 16S data than just what microbes are present and their abundances. Which microbial communities differ and why is the question at the centre of understanding the contribution of the microbiome to human health White et al. (2009), as alterations and changes in microbiomes have been associated with a range of diseases including obesity Turnbaugh et al. (2008, 2009); Duncan et al. (2008), Crohn’s disease Gevers et al. (2014) or ankylosing spondylitis Costello et al. (2015).

A number of statistical analysis tools have been proposed to examine differences between microbial communities as well as identify features that are key to driving the differences. Many tools were originally developed for digital count data (SAGE, RNA-sequencing), such as EdgeR, DESeq, and DESeq2 Robinson and Smyth (2008); Anders and Huber (2010); Love et al. (2014), which model the entity counts from a Negative Binomial distribution. In their thorough simulation study, McMurdie and Holmes (2014) showed that the DESeq2 model performed well in simulated microbiome data compared to other statistical methods developed for RNA-Seq differential abundance analysis. Specific methods were also developed for microbiome data to accommodate for their specific *sparse* nature. Originally, White et al. (2009) proposed Metastat, a non parametric t-test based on permutation or a Fisher’s exact test when data are sparsely sampled. Their approach was a first step towards identifying organisms whose differential abundance correlated with disease. Recently, Paulson et al. (2013) developed a zero-inflated Gaussian (ZIG) distribution mixture model to account for biases due to undersampling of the microbial community.

Another characteristic of microbiome data is their underlying compositional structure due to the varying sampling/sequencing depths between samples from the high-throughput sequencing technologies. After identifying OTU in a sample, it is therefore common to convert each OTU count into relative abundance (proportion) by dividing its count by the total number of counts in each sample. However, as underlined by Aitchinson and colleagues Aitchison (1982), those compositional data reside in a simplex sample space rather than the Euclidian space. Conventional statistical methods such as correlation coefficients Lovell et al. (2015); Ban et al. (2015) or univariate methods may therefore lead to spurious results as the independence assumption between predictor variables is not met. The microbiome data analysis field is showing a growing list of references which mention the limitation of such methods for compositional data Mandal et al. (2015); Fernandes et al. (2014). Therefore, for co-expression analysis, the use of inverse covariance or regularised inverse covariance matrices have recently been proposed for microbiome analysis Kurtz et al. (2015); Ban et al. (2015). Since statistical data analysis is usually carried out in the Euclidean geometry and not in the Aitchison geometry, Aitchison (1982) proposed to transform compositional data into the Euclidian space using centered log ratio (CLR). CLR consists in dividing each sample by the geometric mean of its values and taking the logarithm. Standard univariate and multivariate methods can then be applied on the CLR data Mandal et al. (2015); Kalivodová et al. (2015).

One major drawback of univariate statistical approaches commonly used for 16S rRNA data is that they test each OTU feature individually Robinson and Smyth (2008); Anders and Huber (2010); Paulson et al. (2013), therefore disregarding interactions or correlations between features. On the contrary, multivariate methods treat the entire subset of microbial abundances as a whole, and enable insights into how microbial communities modulate and influence biological pathways. Only a few multivariate approaches have been proposed for 16S analysis, including the Bayesian framework ALDex2 Fernandes et al. (2014) for compositional data, and LEfSe Segata et al. (2011). However those approaches still use univariate tests (Welch’s t‐ or Wilcoxon rank test) to assess the significance of each OTU.

Most multivariate approaches are used to visualise diversity patterns only. Amongst those, Principal Coordinate Analysis (PCoA, a.k.a multidimensional scaling) is an unsupervised approach based on sample-wise distance/dissimilarity matrices such as Bray-Curtis Bray and Curtis (1957), unweighted Lozupone and Knight (2005) or weighted Unifrac Lozupone et al. (2007) to scale for species abundance Gower (1998). Between-class analysis Dolédec and Chessel (1987) is a supervised approach combining PCoA in a supervised framework to segregate sample groups. Those exploratory multivariate approaches give a first insight into similarities between samples, body sites or habitats, but they cannot help identifying key indicator species that discriminate the different groups of samples.

Another critical issue in microbiome data analysis is high inter-subject variability Turnbaugh et al. (2009), which is often reduced with an appropriate experimental repeated-measures design where each subject acts as its own control. Thus, microbial sampling is performed repeatedly on the same subjects over different habitats. Such experimental design has been widely adopted by community profiling studies such as the Human Microbiome Project (HMP, Human Microbiome Project Consortium (2012a,b)) which aims are to define a ‘healthy’ microbiome community by characterising different body sites in the same subjects. However, very few statistical approaches have taken advantage of this design and accommodate for inter-subject variability.

We introduce `mixMC`, a multivariate analysis framework or 16S compositional data to identify OTU features discriminating multiple groups of samples `mixMC` addresses the limitations of existing multivariate methods for microbiome studies and proposes unique analytical capabilities compared to existing statistical methodologies: it can handle repeated-measures experiments and multiclass problems; it provides an internal feature selection procedure to highlight important discriminative features, and friendly interpretable graphical outputs to better understand how microbial communities contribute to each habitat. We applied our approach to multiple body site studies in healthy individuals from HMP and the study from Koren et al. (2011), compared it with existing statistical approaches for microbiome analysis and provide thorough interpretations of the microbial communities unraveled during the multivariate analysis process.

## II. Methods

We analysed publicly available 16S data from the NIH Human Microbiome Project and cross-compared our results with the microbiome study from Koren et al. (2011). The data were processed by the open-source bioinformatics software QIIME Caporaso et al. (2010) for the 16S variable region 1-3. We first describe the different processing, and normalisation steps, and the statistical methods applied in this study, summarised in Figure 1**A**.

**Figure 1:**
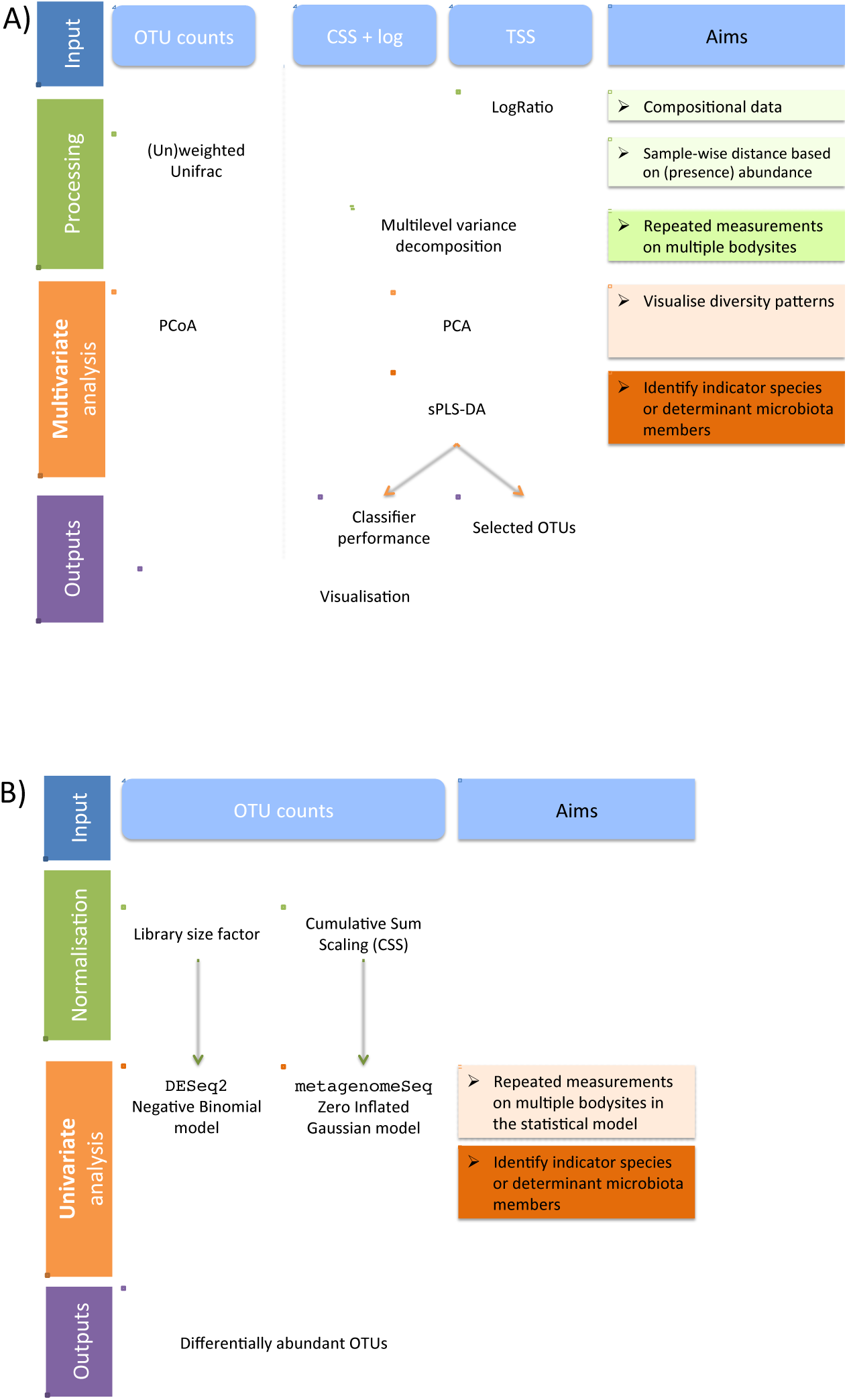
Comparison between multivariate and univariate statistical analysis frameworks for 16S microbiome data. **(A)** Multivariate **`mixMC`** framework including processing/normalisation, optional repeated measures design, unsupervised and supervised analyses, **(B)** Univariate framework, including normalisation and optional repeated measures design analysis

### Data processing and normalisation

One of the characteristics of taxonomic data is that features are often absent from most samples (zero counts), which makes preprocessing and normalisation steps crucial when the aim is to describe microbial communities.

#### Prefiltering

Work by Bokulich et al. (2013) demonstrated that strict quality filtering of reads greatly improves taxonomy assignment and alpha diversity measures for microbial community profiling. After removing samples with a very low number of total OTU counts (less than 10), our prefiltering step consisted in removing lowly abundant OTU as these OTU may not be informative to fully characterise microbial environments. OTU were removed if their average abundance across all samples was below 0.01%. While this prefiltering step may result in a very small number of remaining OTU to analyse, it avoids spurious results in the downstream statistical analysis and counteracts sequencing error. The threshold is the default value proposed in QIIME and also used in other microbiome studies (Knights et al. (2011); Arumugam et al. (2011) to cite a few).

#### Normalisation

The issue of sparse counts also needs to be accounted for during normalisation Paulson et al. (2013). Thus, the normalisation technique needs to be chosen carefully as it can have strong repercussions in the statistical results. So far, two types of normalisations have been proposed for microbiome studies.

The Total Sum Scaling normalisation (TSS) is the most commonly used approach which divides each OTU count by the total number of counts in each individual sample to account for uneven sequencing depths across samples. However, this normalisation measures relative information (i.e. proportions) and results in data residing in a simplex rather than the Euclidian space Aitchison (1982). Therefore, statistical methods are not directly applicable on TSS data and can lead to spurious false discoveries. The solution is to transform TSS data to project them to an Euclidian space using log ratio transformations. The Centered Log Ratio transformation (CLR) has been recently applied in several compositional data studies Fernandes et al. (2014); Mandal et al. (2015); Kalivodová et al. (2015). Let *x* = (*x*_1_,…, *x*_*p*_)′ denote a composition on the *p* TSS normalised OTU counts, then the CLR transformation is defined as

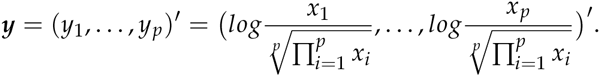

In **`mixMC`** CLR transformation is applied on TSS normalised data (Fig. 1**A**).

The Cumulative Sum Scaling normalisation (CSS, Paulson et al. (2013)) has been developed to prevent TSS bias in differential abundance when few measurements are sampled preferentially. CSS can be considered as an extension of the quantile normalisation approach and consists ofTSS scaling raw counts that are relatively invariant across samples, up to a percentile determined using a data-driven approach. Therefore, CSS combines both TSS and quantile-type normalisations. According to the authors, CSS partially accounts for compositional data. Consequently, and as proposed in the metagenomeSeq package Paulson et al. (2015) the raw OTU counts were log transformed and CSS normalised prior to the statistical analysis in our framework (Fig. 1**A**).

### Methods

One of the main objectives of our study is to extend and apply multivariate statistical analysis methods for microbiome compositional data. The **`mixMC`** framework (Fig. 1**A**) includes unsu-pervised analyses to visualise diversity patterns with Principal Component Analysis (PCA) and supervised analyses to identify indicator species or determinant microbiota members characterising differences between habitats or body sites (sparse Partial Least Square Discriminant Analysis, sPLS-DA). In addition, our framework addresses a commonly encountered experimental design called *repeated-measures design,* where microbial sampling is performed on the same individuals but in different body sites to detect differences between habitats. This design leads to analytical challenges in order to be able to discern subtle differences between body sites from the large variation between individuals within the same body site.

#### Unsupervised multivariate analysis and ILR transformation

PCA variants, such as Principal Coordinate Analysis (PCoA, a.k.a multidimensional scaling) allows for dimension reduction of the data and visualisation of diversity patterns in microbiome study. PCoA is commonly applied to non Euclidian sample-wise dissimilarity matrices (e.g. Bray-Curtis) or phylogenetic distances between sets of taxa in a phylogenetic tree (weighted or unweighted Unifrac distance, Lozupone and Knight (2005); Lozupone et al. (2007)). Alternatively, PCA can be applied on log ratio compositional data using the Isometric Log Ratio transformation (ILR). According to Filzmoser et al. (2009), the ILR transformation is preferable to the CLR transformation that may result in singular variance-covariance matrix. ILR transformed data (*z*_1_,&, *z*_*p*–1_)′ are spanned by (*p* – 1) new coordinates with respect to the orthonormal basis *V*, such that

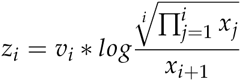

with the orthonormal basis vector 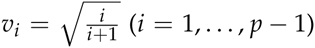 Egozcue et al. (2003). However, ILR transformed data are not easily interpretable as they are of dimension *p* – 1, i.e. there is not a one-to-one transformation of all features. Therefore, Filzmoser et al. (2009) proposed to back transform the PCA results to the CLR space using the linear relationship between CLR and ILR transformations: *y* = *Vz*, where *V* = (*v*_1_,…, *v*_*p* – 1_) is a *p* × (*p* – 1) matrix with orthonormal basis vectors *v*_*i*_ as defined above. We applied PCA on ILR transformed data using customised R scripts from the robCompositions package Templ et al. (2011).

#### Multilevel variance decomposition

One way to account for repeated measurements designs is to separate the body site variation (termed ‘*within variation*’) from the individual variation (termed ‘*between subject variation*’) via variance decomposition. In univariate analyses, this step refers to repeated measures ANOVA (also called within-subjects ANOVA). In multivariate analysis we refer to a multilevel approach as proposed by Westerhuis et al. (2010). The within subject variation is obtained by calculating the net differences between repeated observations, i.e. between each body site within each individual. Since the within subject variation assesses the difference in the body sites within each subject and disregards the possibly large individual variation, the within variation can then be used as input data in the subsequent multivariate statistical analysis as was proposed by Liquet et al. (2012). In **`mixMC`**, the multilevel variance decomposition is applied on the log ratio transformed data described above, prior to the multivariate analyses (Fig. 1**A**). Note that the variance decomposition in the multilevel approach does not take into account the correlation structure or order between the measurements and is not appropriate for a time course experiment where the objective is to examine the effect of time in a study (see for example applications of linear mixed model splines for those cases Straube et al. (2015); Paulson et al. (2015)).

#### Supervised multivariate analysis

The multivariate approach sparse Partial Least Squares Discriminant Analysis (sPLS-DA, Lê Cao et al. (2011)) is an extension of the PLS algorithm from Wold et al. (2001) to perform feature selection with multilevel decomposition Liquet et al. (2012). In **`mixMC`** we further extended the multilevel sPLS-DA for microbiome data using either CSS normalised data, or CLR transformed TSS data. For the latter, we restricted the log transformation to CLR as the sPLS-DA framework can only be applied to *p* dimensional data to identify and select indicator species internally in the statistical model.

##### Principle of PLS-DA

PLS-Discriminant Analysis is a multivariate regression model which maximises the covariance between linear combinations of the OTU counts and the outcome (a dummy matrix indicating the body sites for each sample).Covariance maximisation is achieved in a sequential manner via the use of latent component scores. Each component is a linear combination of OTU counts and characterises a particular source of co-variation between the OTU and the body sites. As a consequence, the final number of components summarising most of the information from the data must be specified. The sparse version of PLS-DA uses Lasso penalisation Tibshirani (1996) to select the most discriminative features in the PLS-DA model. The penalisation is applied componentwise and the resulting selected features reflect the particular source of covariance in the data highlighted by each PLS component Lê Cao et al. (2011).

##### Parameters and performance evaluation

The number of features to select per component must be specified in sPLS-DA and is usually optimised using cross-validation.In this study we used 10-fold cross-validation repeated 100 times. For varying features selected by the sPLS-DA model, the classification error rate resulting from the cross-validation process was then recorded. The optimal number of selected features was chosen so as to achieve the lowest error rate on each component. This procedure was concurrently used to choose the total number of components. Once these parameters were chosen, the final sPLS-DA model was run on all samples to obtain the final lists of discriminative OTU features on each component.

##### Graphical and numerical outputs

We further characterised each selected OTU by calculating its median normalised count in each body site. An OTU was defined as ‘contributing to a body site’ if the median count in that specific body site is higher than in any other body site. We graphically represented the contribution of each selected OTU with a barplot where each OTU bar length corresponds to the importance of the feature in the multivariate model (i.e. the multivariate regression coefficient with either a positive or negative sign of that particular feature on each component) and the colour corresponds to the body site in which it was the most abundant. The contribution plot displays bacterial taxonomy at the family level. Another type of graphical output used GraPhlAn Asnicar et al. (2015) to produce circular representations of taxonomic trees. The results from sPLS-DA were directly included in GraPhlAn to complement the contribution plot with taxonomy information. Other insightful outputs included sample plots where each individual is projected on the sPLS-DA components, the list of OTU features selected on each component, the cross-validation error rate per component and the number of features contributing to each body site and each component.

The multilevel sPLS-DA framework is implemented in the R package **mixOmics** Lê Cao et al. using the multilevel module Liquet et al. (2012). The cladogram was generated using the GraPhlAn Python code Asnicar et al. (2015). All R codes and tutorials are available on our website www.mixOmics.org/mixMC

#### Univariate analysis

We considered univariate statistical approaches capable of handling repeated-measures experiments. Unlike the supervised multivariate approach presented above, the univariate methods output a p-value for each OTU testing for differential abundance between body sites. P-values were adjusted for multiple testing using the False Discovery rate Benjamini and Hochberg (1995) at a significance level of 5%. We considered two univariate approaches, DESeq2 and ZIG (Fig. 1 **B**).

DESeq2 was developed for DNA sequencing read count data where mean and variance for the binomial distribution is estimated for each feature Anders and Huber (2010).The OTU counts are normalised internally to the method with respect to a library size factor estimation. DESeq2 can be adapted for microbiome analysis and often serves as a basis of comparison to novel methodological developments Paulson et al. (2013); McMurdie and Holmes (2014); Fernandes et al. (2014). We compared DESeq2 differentially abundant OTU to those selected with `mixMC`. However, the reader must keep in mind that the normalisation in DESeq2 does not address the issue of compositional data, but the generalised linear model interface in DESeq2 enables repeated measurements experimental designs. We used mean dispersion estimates as implemented in the R package `DESeq2` Love et al. (2014).

ZIG is a mixture model with a Zero-Inflated Gaussian distribution that was recently proposed by Paulson et al (2013) to account for varying depths of coverage that is typical for microbial community undersampling. In the ZIG model, OTU counts are first log transformed and then CSS normalised.

ZIG uses a linear model framework which can include a repeated-measures design and is implemented in the R package metagenomeSeq Paulson et al. (2015).

### Case studies

#### HMP case studies

We analysed subsets of the NIH HMP16S data downloaded from http://hmpdacc.org/HMQCP/all/ for the V1-3 variable region. The original data contained 43 146 OTU counts for 2 911 samples measured from 18 different body sites. We focused on the first visit of each healthy individual and further divided the data into two data subsets. For both data sets a preliminary exploratory PCoA confirmed that there was no confounding covariate effect due to run center or gender (see Suppl. Material S4).

##### Most diverse body sites dataset

Understanding microbial community diversity across body habitats is fundamental to study the human microbiome. In their extensive HMP data statistical analysis, Li et al. (2012) quantified intra-sample diversity based on Shannon index. Based on their results we chose three main types of habitats which were the most diverse in terms of all genera-based and OTU-based taxonomic units (Tables 1 and 2 in Li et al. (2012)). Those body sites were Subgingival plaque (Oral), Antecubital fossa (Skin) and Stool sampled from 54 unique individuals for a total of 162 samples. The prefiltered dataset included 1 674 OTU counts (Table S1). *Oral body sites dataset.* While many published analyses have focused on the main microbial habitats (gut, oral cavity, skin and vagina from the Human Microbiome Project Consortium (2012b,c)), little has been done to comprehensively characterise multiple sites within a single habitat. In this data set we solely considered samples from oral cavity, which has been found to be as diverse as the stool microbiome Li et al. (2012). The nine oral sites were Attached Keratinising Gingiva, Buccal Mucosa, Hard Palate, Palatine Tonsils, Saliva, Subgingival Plaque, Supragingival Plaque, Throat and Tongue Dorsum. After prefiltering, the data included 1 562 OTU for 73 unique patients and a total of 657 sample (Table S1).

**Table 1.** Oral data. Top: Number of selected features at the OTU (family) level and mean classification error rate per component. Bottom: Number of features at the OTU (family) contributing to each body site for each sPLS-DA component. An OTU is defined as contributing to a body site if the median count in that specific site is higher than in any other site. The median counts were calculated from the multilevel normalised data. Note that we can observe some overlap between families across the different body sites.

|  | Comp 1 | Comp 2 | Comp 3 | Comp 4 | Comp 5 | Comp 6 | Comp 7 | Comp 8 |
| --- | --- | --- | --- | --- | --- | --- | --- | --- |
| # features selected | 60 (13) | 40 (2) | 190 (18) | 200 (14) | 40 (8) | 200 (26) | 180 (23) | 190 (22) |
| mean classification error rate | 0.778 | 0.584 | 0.501 | 0.410 | 0.336 | 0.316 | 0.279 | 0.262 |
| sd classification error rate | 0.000 | 0.002 | 0.003 | 0.003 | 0.003 | 0.005 | 0.004 | 0.004 |
| Attached Keratinized gingiva | 0 | 35 (2) | 123 (12) | 9 (6) | 1 (1) | 73 (16) | 34 (11) | 47 (15) |
| Buccal mucosa | 0 | 5 (1) | 4 (1) | 1 (1) | 0 | 31 (4) | 3 (1) | 3 (1) |
| Hard palate | 2 (1) | 0 | 1 (1) | 3 (1) | 0 | 3 (2) | 5 (3) | 9 (3) |
| Palatine Tonsils | 1 (1) | 0 | 0 | 5 (3) | 0 | 2 (2) | 4 (2) | 6 (4) |
| Saliva | 5 (3) | 0 | 2 (2) | 28 (5) | 0 | 4 (2) | 11 (5) | 7 (2) |
| Subgingival plaque | 0 | 0 | 7 (7) | 15 (5) | 39 (7) | 14 (11) | 6 (5) | 21 (10) |
| Supragingival plaque | 11 (4) | 0 | 53 (8) | 23 (6) | 0 | 31 (9) | 15 (8) | 31 (6) |
| Throat | 11 (5) | 0 | 0 | 16 (4) | 0 | 5 (3) | 42 (5) | 9 (4) |
| Tongue dorsum | 30 (9) | 0 | 0 | 100 (8) | 0 | 37 (11) | 60 (13) | 57 (12) |

#### Koren dataset

Koren and colleague Koren et al. (2011) examined the link between oral, gut and plaque microbial communities in patients with atherosclerosis and controls. To compare our results with the HMP most diverse study, only the healthy individuals were retained in the analysis. This study contained partially repeated measures from multiple sites including 15 unique patients samples from saliva and stool, and 13 unique patients only sampled from arterial plaque samples. The data were downloaded from the Qiita database (http://qiita.microbio.me/study/description/349) and included 5 138 OTU. After prefiltering, the data included 973 OTU for 43 samples.

## III. Results

### Unsupervised analyses on Most Diverse body sites dataset

Unsupervised analyses such as PCoA or PCA on ILR transformed data are useful to visualise diversity patterns between microbial communities. We compared the PCoA and PCA sample visualisation for different types of normalisations (TSS-ILR, CSS) followed by a multilevel variance decomposition for repeated measures.

#### PCoA

A PCoA performed on the filtered OTU counts (with no normalisation) showed that the unweighted Unifrac distance could highlight diversity patterns between each body site better than weighted Unifrac (Fig. 2). As this study focuses on the most diverse body site, the presence or absence of microbial communities is to be expected. Applying PCoA on the unfiltered count data led to the same interpretation (Suppl. Fig. S1), but we observed a lower amount of explained variance of the first and second coordinate as more ‘noisy’ OTU are present in the data (unweighted Unifrac: 11.28% and 8.95% for the unfiltered data vs. 17.37% and 14.48% for the filtered data).

**Figure 2:**
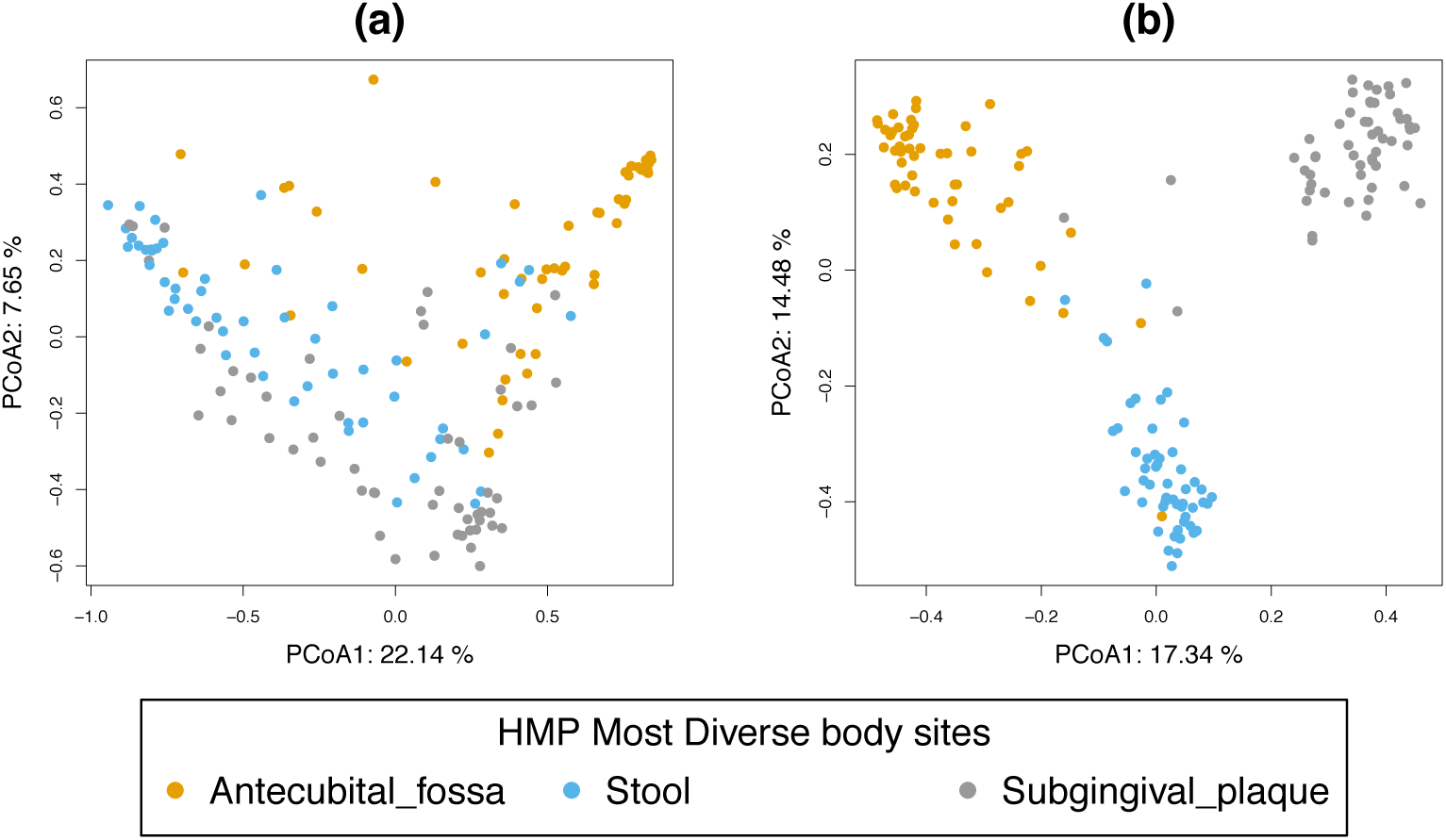
Most diverse data, PCoA sample plots. Sample plot on the first two coordinates with (a) weighted Unifrac (b) unweighted Unifrac calculated on the filtered OTU count table (based on 1,674 OTU).

#### PCA

We then assessed the effect of the different normalisation strategies as well as the multilevel variance decomposition. Each type of transformation (TSS, TSS + ILR OTU count, CSS) seemed to cluster samples similarly according to the body sites (Fig. 3 **(a) (c) (e)**). The differences arose when we added the multilevel decomposition, leading to a smaller variability within body sites and a greater variability between body sites (Fig. 3 **(b) (d) (f)**). Consequently, the explained variance per component was larger than previously observed. The use of the normalisations TSS-ILR or CSS also increased the explained variance, with a maximal cumulative explained variance attained with TSS-ILR of 44.6% for the first two components (33.5% for CSS).

**Figure 3:**
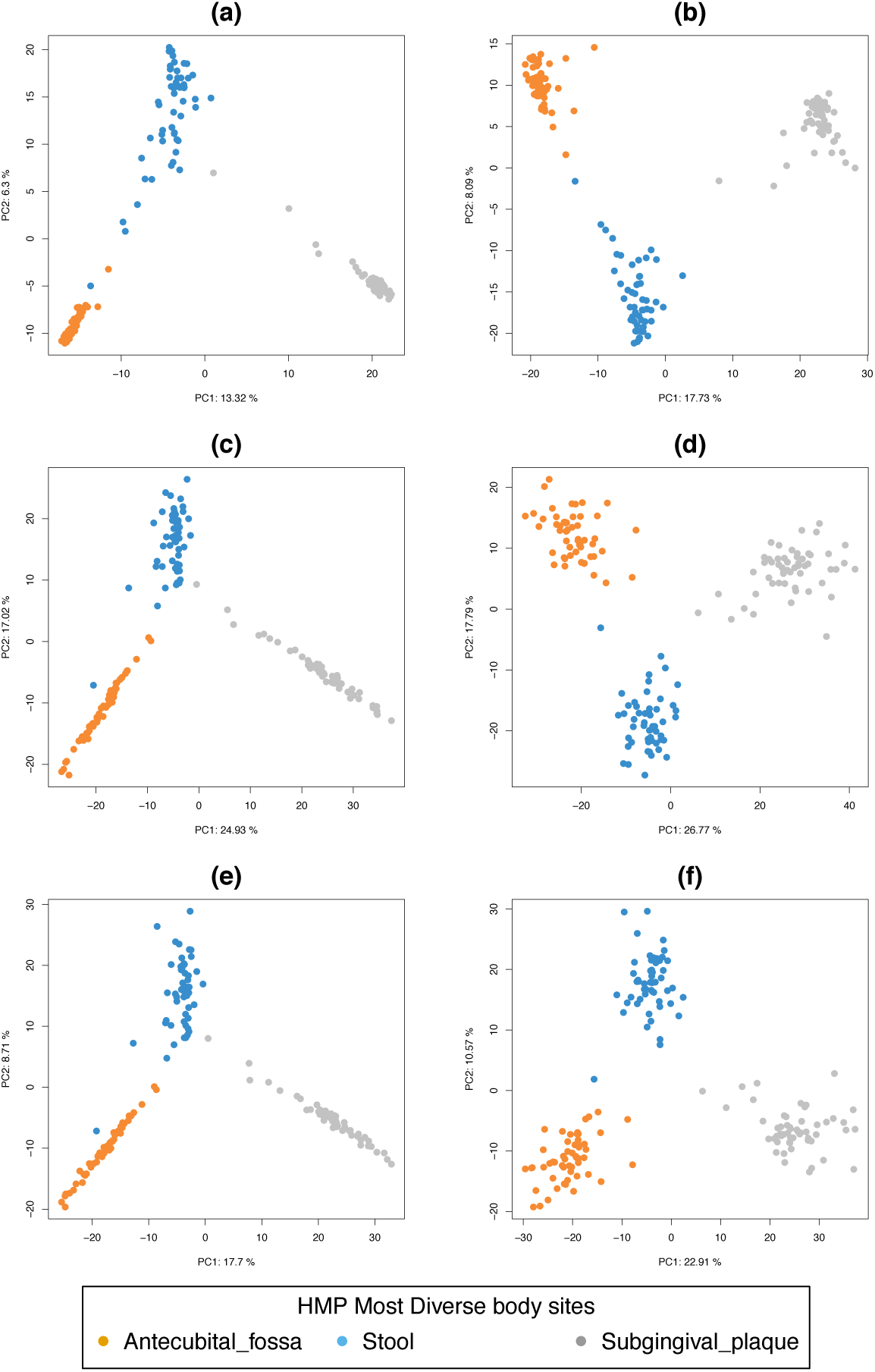
Most diverse data, PCA sample plots. (a) TSS and (b) TSS multilevel OTU log counts, (c) TSS-ILR and (d) TSS-ILR multilevel normalised counts, (e) CSS and (f) CSS multilevel log counts. Colours indicate body sites and the percentage of explained variance for each principal component is indicated in the axes labels.

### Supervised analysis on Most Diverse body sites dataset and OTU selection

Our preliminary exploration using unsupervised multivariate analyses indicated that the abundance of microbial communities could characterise each body site quite clearly. The multilevel decomposition enabled better separation of the body site clusters, in particular when applied to the TSS-ILR or CSS normalised data.

The next step was to perform a supervised analysis with multilevel sPLS-DA in order to identify a microbiome signature characterising each body site. We compared the different normalisation strategies (TSS-CLR or CSS) in our multivariate method to DESeq2 and ZIG univariate methods.

#### The impact of normalization to identify discriminative features with sPLS-DA

The sPLS-DA classification performance was similar in both TSS-CLR or CSS normalised data, and a minimum classification error rate was reached for two components in the multivariate model (0.7% for TSS-CLR and 0.3 % for CSS, Table S4). For both normalizations the first component consistently misclassified antecubical fossa on the first component but correctly classified the two other body sites. The addition of the second component enabled a better classification of all body sites (Fig. 4). The optimal number of selected OTU and number of components were chosen so as to achieve a lowest overall classification error rate componentwise using 10-fold cross-validation repeated 100 times. The final selected list include 160 (130) OTU with TSS-CLR (CSS), see Table S5. Since the sPLS-DA is fitted in a sequential manner with one component at a time, we can also assess the contribution of the selected OTU selected on each component (Table S5).

**Figure 4:**
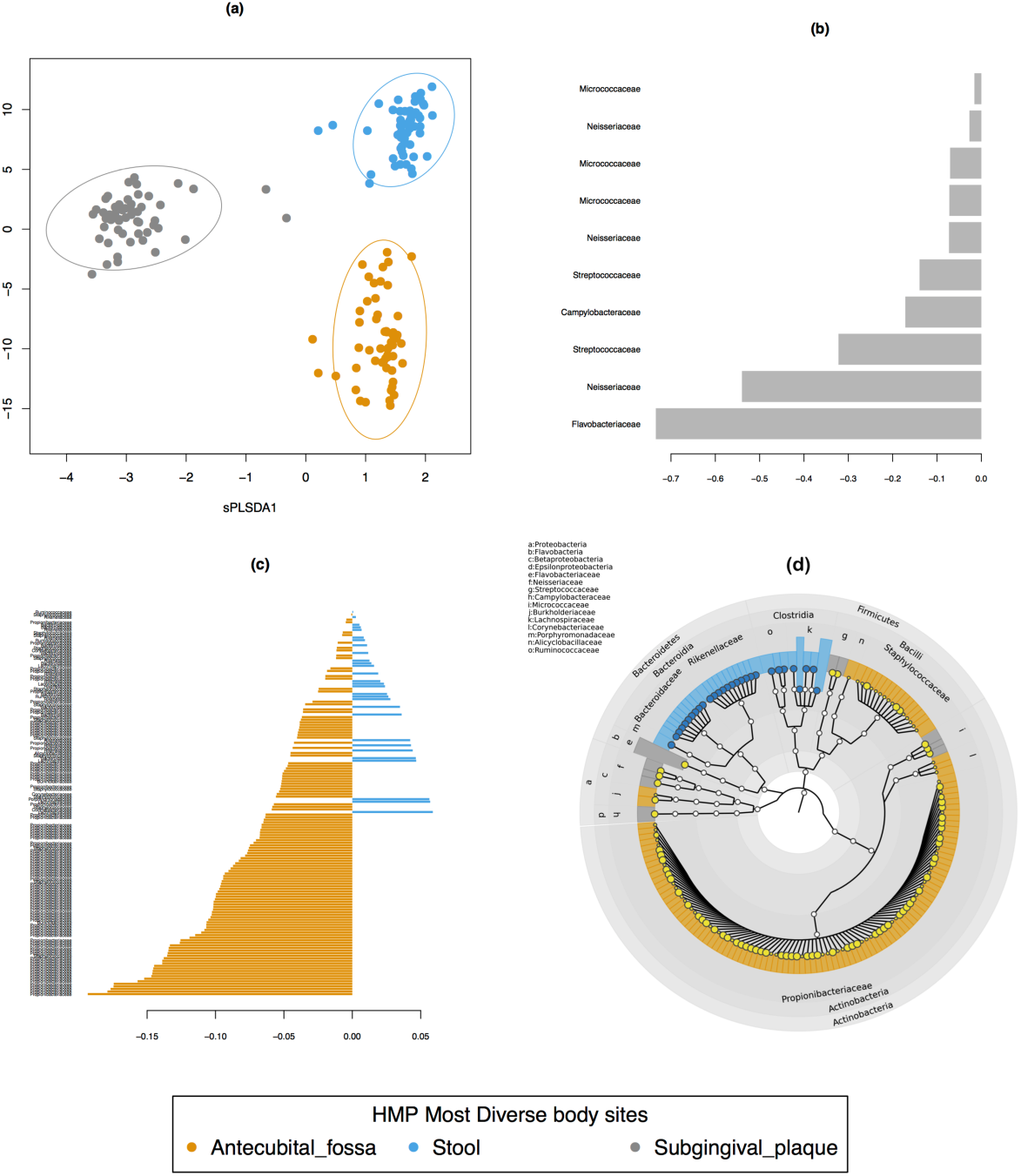
Most diverse TSS-CLR data, sPLS-DA sample, contribution and cladogram plots. (a) sample plot on the first two components with 95% confidence level ellipse plots, (b) and (c) represent the contribution of each OTU feature selected on the first (10) and second component (120) respectively, with OTU contribution ranked from bottom (important) to top. Colours in the contribution plot indicate the body site with the highest median for each selected OTU labelled at the family level. The negative (resp. positive) sign on the x-axis represents the regression coefficient weight of each feature in the linear combination of the sPLS-DA component. (d) Cladogram generated from the sPLS-DA result using GraphlAn: background colour indicates the body sites where the OTU is most present, the node size represent the median OTU count for that contributing body site and the node colour indicates a negative (black) or positive (yellow) weight from the sPLS-DA weight vector as shown in (b) and (c).

We found that both normalizations identified very similar bacterial families. Component 1 was found to characterise the subgingival plaque including *Micrococcaceae, Neisseriaceae, Streptococcaceae, Flavobacteriaceae* and *Campylobacteraceae*. In addition, CSS also identified the *Burkholderiaceae* family. Component 2 characterised stool and anticubital fossa. For the latter body site, TSS+CLR normalization identified *Propionibacteriaceae*, *Staphylococcaceae and Corynebacteriaceae* while CSS also identified *Propionibacteriaceae*, *Staphylococcaceae* but failed to identify *Corynebacteriaceae.* Bacterial families characterising stool included *Bacteroidaceae*, *Ruminococcaceae*, *Lachnospiraceae*, *Rikenellaceae* and *Porphyromonadaceae*. Across the three body sites, we found that both normalizations led to very similar families of bacteria - 5 families for component 1, 10 (TSS-CLR) or 8 (CSS) for component 2 with a difference of 1 or 2 families on each component between TSS-CLR and CSS, see Table S5. Interestingly, we observed that increasing the number of selected OTU did not add more relevant bacteria families for that CSS normalisation. Rather, the proportion of number of OTU corresponding to the families varied (Fig. 4 **(d)**).

#### Comparisons with no multilevel approach

In order to understand the impact and benefits of the proposed multilevel approach, we examined the OTU selected by sPLS-DA multilevel on either the TSS or CSS normalised counts without multilevel transformation. The classification error rate was substantially greater than with the previous multilevel analysis, (6% for TSS-CLR and 3% for CCS for two components) with a larger number of OTU selected (400 OTU selected for TSS-CLR and 240 for CSS).

With the TSS-CLR normalisation, we identified similar families characterising subgingival plaque on the first component, including *Burkholderiaceae*, *Fusobacteriaceae*, *Gemellaceae*, *Veillonel-laceae* The families selected on the second component characterised antecubital fossa similarly to the multilevel approach, however the notable omission was the entire *Ruminococcus* family characterising stool in the multilevel approach that was not identified in the non multilevel approach. These observations led us to the conclusion that a classical multivariate analysis ignoring the repeated-measures design tended to identify differential features driving the overall signature and disregarded subtleties between microbial communities in environments sampled on the same individuals.

#### Comparison with univariate analysis

We then compared the univariate approaches DESeq2 and ZIG for assessing differential abundance with a repeated measures design. The total number of OTU selected differed slightly between those two approaches (Supp. Table S2). We compared the differentially abundant OTU selected by either DESeq2 or ZIG to the discriminative OTU selected by sPLS-DA with either TSS-CLR or CSS normalization (Supp. Fig. S2). We observed strong differences between the univariate approaches at both OTU and family levels. Interestingly, the sPLS-DA selections were all included in the ZIG and DESEq2 selections (Fig. S2).

DESeq2 identified relevant features in common with sPLS-DA selections, such as *Propionibacteriaceae*, *Staphylococcaceae* and *Corynebacteriaceae* with the addition of *Burkholderiaceae* as a defining feature characterising Antecubital fossa. It also characterised the Subgingival plaque microbial community with OTU from *Streptococcaceae*, *Neisseriaceae*, *Gemellaceae* and *Micrococcaceae* families, which were also identified in the sPLS-DA analyses. The only drawback of DESeq2 was the lack of Stool characterization. Indeed, very few bacterial families, including Bacteroides and *Lachnospiraceae*, were identified. Such low bacterial diversity was not consistent with the sPLS-DA nor with the literature. Similar to DESeq2 and sPLS-DA, ZIG identified features of the Antecubital fossa with OTU belonging to *Propionibacteriaceae*, *Staphylococcaceae*, *Burkholderiaceae* and *Corynebacteriaceae*. Like DESeq2, ZIG described the Subgingival plaque microbiome with OTU belonging to *Streptococcaceae, Neisseriaceae, Micrococcaceae* and *Gemellaceae*. However, the ZIG analysis also identified OTU belonging to *Fusobacteriaceae*, *Burkholderiaceae*, *Flavobacteriaceae*, *Campylobacteraceae*, *Veillonellaceae* and *Actinomycetaceae*. In contrast to DESeq2, ZIG identified and described the Stool microbiome well, with OTU belonging to the families of *Bacteroidaceae*, *Porphyromonadaceae*, *Rikenellaceae*, *Lachnospiraceae* and *Ruminococcaceae.*

### Analysis of the oral body site dataset with mixMC

Similar to the Most Diverse data set, unsupervised data analyses showed that unweighted Unifrac better discriminated the different body sites (plaque, gingiva) compared to weighted Unifrac in the PCoA sample plots (Fig S3 **(a-b)**). When comparing TSS-ILR with CSS normalised counts, TSS-ILR explained greater variance (21.35% on the first component) than CSS (13.63%), with clearer clusters corresponding to the body sites (Fig. S3 **(c), (e)**). The explained variance further increased with a multilevel variance decomposition (25.37% vs. 18.22%, Fig. S3 **(d), (f)**).

#### sPLS-DA performance and choice of parameters

We observed similar classification performances between sPLS-DA on either TSS-CLR or CSS, with a slightly lower classification error rate for TSS-CLR (Fig. S5). sPLS-DA is a model that builds on successive components that are progressively added. For each component, an optimal list of OTU features was selected using-cross validation (Table 1). The final sPLS-DA model included 8 components that led to optimal performance, with a classification error rate that substantially decreased from 78% (component 1) to 26% for TSS-CLR and 30% for CSS (component 8). The classification error rate was still quite high as similar body sites were consistently misclassified across components (Table S3). For example, Tonsils retained the highest error rate as no OTU was able to characterise that particular body site (Table 1).

We observed that the TSS-CLR normalisation was better at characterising tonsil and plaque (component 1), buccal mucosa (component 2) and gingiva (component 3) than the CSS normalisation. The CSS normalisation also led to a substantial number of ties when assessing the body site contribution of the selected OTU (not shown). Therefore, the detailed analysis that follows solely focuses on a multilevel sPLS-DA model with TSS-CLR normalisation.

#### Body sites characterisation

Figure 5 displays the sPLS-DA sample representations for the first three components (see Figure S6 for the remaining 5 components). Each of these components seemed to characterise specific subsets of the body sites. For example component 1 discriminated sub and supra gingival plaque against the other body sites, component 2 clustered attached keratinised gingiva and buccal mucosa, but with no clear cut separations from the body sites (Fig. 5 **(a)**), while component 3 seemed to separate attached keratinised gingiva (Fig. 5 **(c)**). Supplemental Figure S6 shows that similar conclusions could be drawn for the other components. The interpretation of these sample plots can be subjective, however, they reflect the close anatomical proximity of the different sample sites in the mouth, such as the tongue coming in contact with the hard palate, teeth, saliva and gums. More insight on the microbial communities can be gleaned as sPLS-DA selects specific sets of features which are linearly combined to determine each component. Given the proximity of some of these body sites, we expect a substantial overlap in some of the bacterial families represented, but we also observed interesting differences which are presented below.

**Figure 5:**
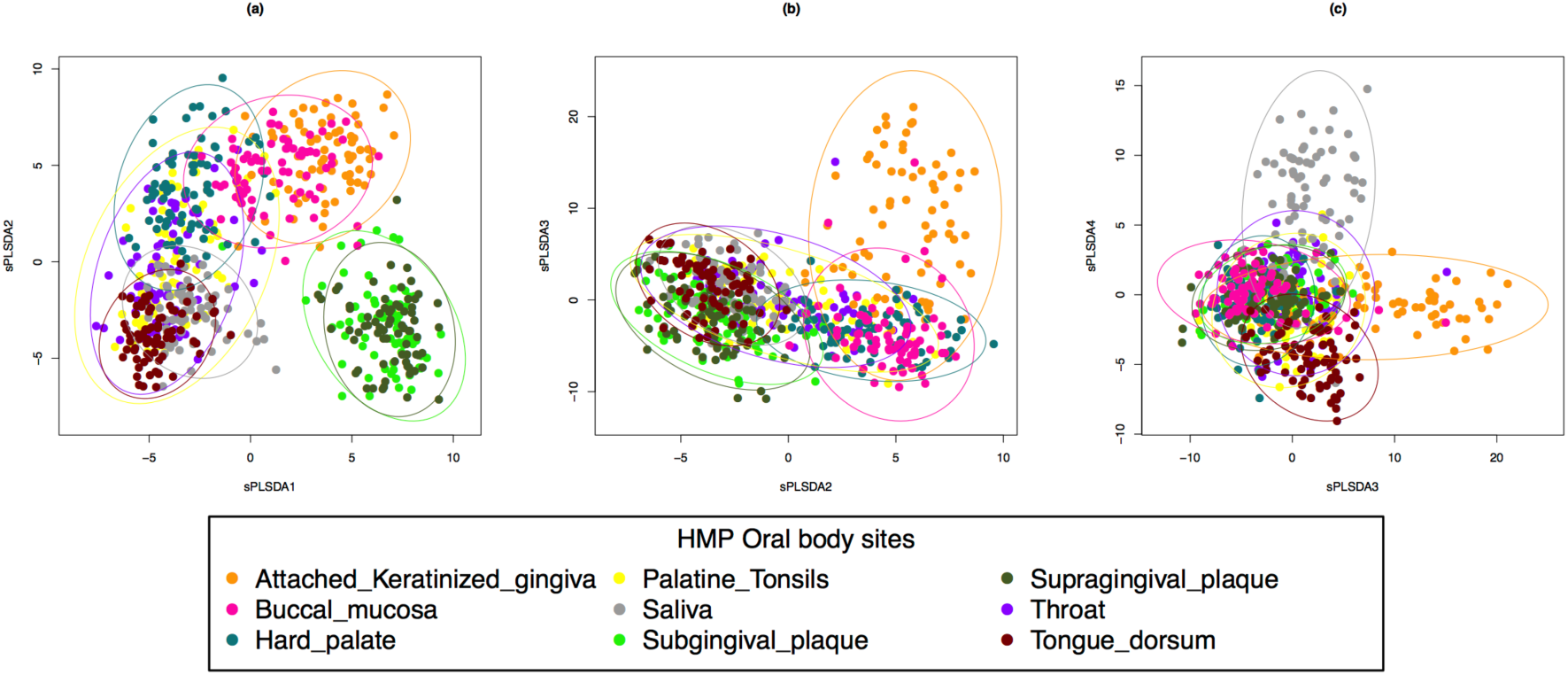
Oral data, sPLS-DA sample plot for the different sPLS-DA components. (a) Component 1 vs. Component 2, (b) Component 2 vs Component 3.Colours indicate the different body sites, 95% confidence ellipses are displayed.

#### Features contribution

In addition to Figure 6, Table 1 shows the number of features contributing to each oral site per component at the OTU and family levels. Those outputs combined with the interpretation from the sample plots in Fig. 5 enable a better insight into bacteria contributing to body sites that are contiguous. For some cases we observed similar contributions of microbial communities in close body sites, for example Throat and Tongue appeared to be characterised by the same family of bacteria. The closeness of those selected bacteria in terms of their taxonomy can be visualised in the cladogram in Figure 6 **(d)**.

**Figure 6:**
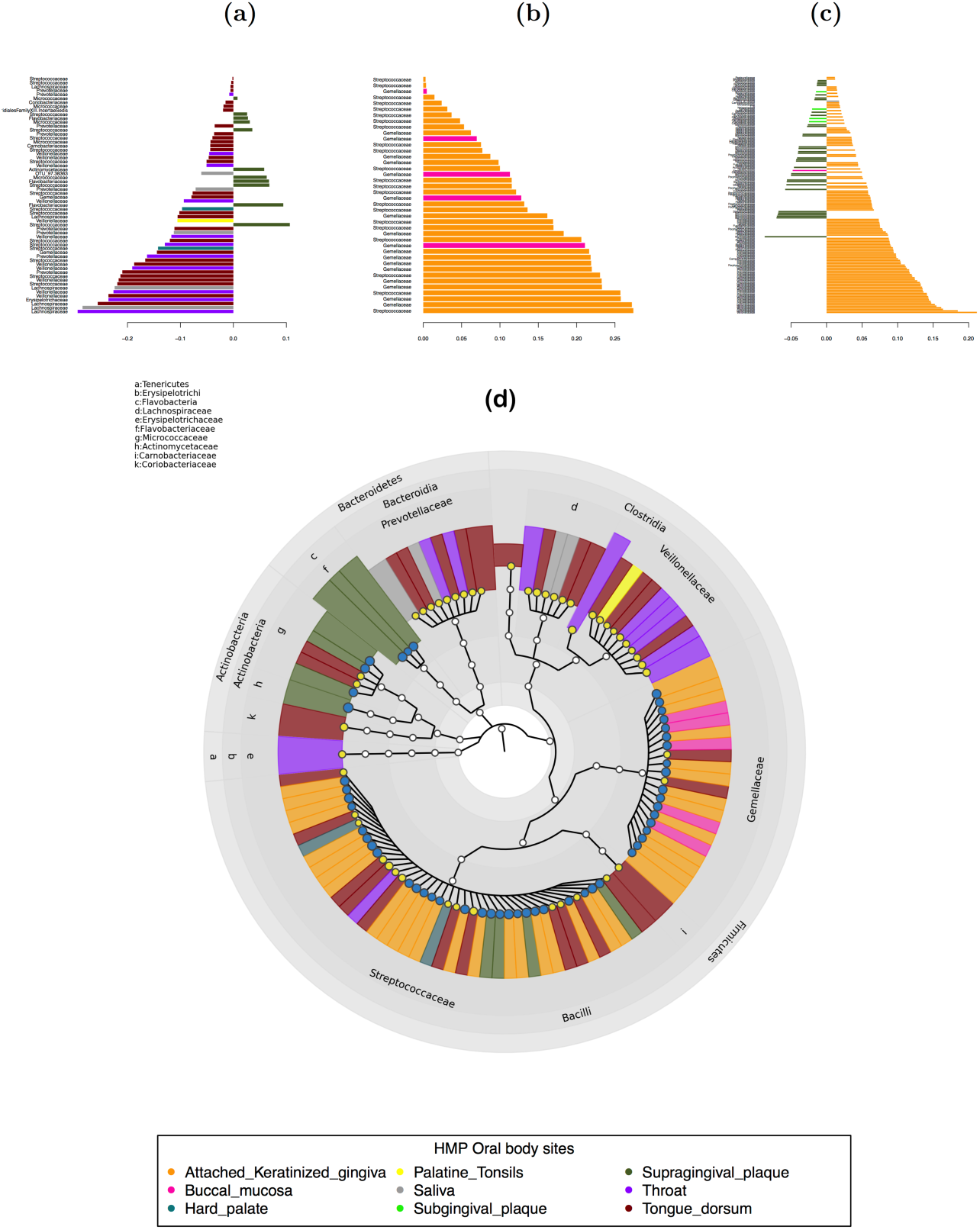
Oral data, contribution and cladogram plots of the features selected for each sPLS-DA component. (a) Component 1, (b) Component 2, (c) Component 3. Colours in the contribution plot indicate the body site with the highest median for each selected OTU labelled at the family level. The negative (resp. positive) sign on the x-axis represents the regression coefficient weight of each feature in the linear combination of the sPLS-DA component. In (c) only the top 150 OTU are represented. (d) Cladogram generated from the sPLS-DA results for components 1 and 2 using GraphlAn: background colour indicates the body sites where the OTU is most present, the node size represents the median OTU count for that contributing body site and the node colour indicates a negative (black) or positive (yellow) weight from the sPLS-DA weight vector as shown in (a) and (b).

We examined the ability of sPLS-DA to highlight subtle differences and characterise different sites in close proximity within the oral microbiome. We list below the relevant families selected on the first three sPLS-DA components, which appear to characterise particular body sites (Table 1, III).

The bacteria families selected on component 1 strongly characterised hard palate (members of the *Streptococcaceae* family), saliva (*Prevotellaceae*, *Lachnospiraceae* as well as the phylum TM7 recently described in He et al. (2015) and found prevalent in oral cavity), supragingival plaque as well as throat and tongue. The throat microbiome was characterized by *Prevotellaceae*, *Lachnospiraceae*, *Veillonellaceae*, *Streptococcaceae* and *Erysipelotrichaceae.* The tongue was found to be more diverse with eight families of bacteria found to be characterising the site. These include the order *Clostridiales* families *Coriobacteriaceae*, *Gemellaceae*, *Carnobacteriaceae*, *Lachnospiraceae*, *Prevotellaceae*, *Micrococcaceae*, *Streptococcaceae* and *Veillonellaceae.* Component 2 was able to separate attached keratinized gingiva from buccal mucosa with the families *Gemellaceae* and *Streptococcaceae.* Component 3 discriminated multiple sites, in particular attached keratinized gingiva (*Prevotellaceae, Porphyromonadaceae, Flavobacteriaceae, Carnobacteriaceae, Streptococcaceae, Fusobacteriaceae, Campylobacteraceae, Pasteurellaceae, Neisseriaceae, Moraxellaceae* and TM7), buccal mucosa and hard palate (*Streptococcaceae* for both). Interestingly, component 3 was able to discriminate subgingival plaque (*Burkholderiaceae, Flavobacteriaceae, Gemellaceae, Micrococcaceae, Neisseriaceae, Prevotellaceae* and *Streptococcaceae*) from supragingival plaque (*Actinomycetaceae*, *Burkholderiaceae*, *Flavobacteriaceae*, *Fusobacteriaceae*, *Micrococcaceae*, *Neisseriaceae* and *Streptococcaceae*) with some overlap between the families.

Applying the **`mixMC`** framework on the oral case study, demonstrated that the resulting graphical and numerical outputs help identifying relevant bacteria families characterising subtle differences in the oral environment while deciphering particular characteristics in each body site.

### Comparison with the Koren data set

⃘ further validate the relevance of our multivariate method to discriminate and identify microbial features describing microbial communities, we applied our sPLS-DA to the study from Koren et al.Koren et al. (2011). Since the dataset only contained partially repeated measures from multiple sites (individual patients samples in plaque were not sampled in other body sites), we applied a non multilevel sPLS-DA on the TSS-CLR data, resulting in a selection of 30+100 OTU on two components (Fig. S7, III). We found that sPLS-DA was able to clearly and distinctly discriminate the three body sites saliva, plaque and stool Component 1 best characterised stool identifying families of bacteria such as *Lachnospiraceae*, *Ruminococcaceae* and *Bacteroidaceae;* similar to what was observed in the HMP dataset. Component 2 best discriminated arterial plaque and saliva. Arterial plaque was characterised by families including *Burkholderiaceae*, *Propionibacteriaceae*, *Pseudomonadaceae and Staphylococcaceae*, which was consistent with what the authors reported to as the ‘core microbiome’ for arterial plaque samples. Our analysis also identified *Alcaligenaceae*, *Enterobacteriaceae*, *Moraxellaceae* and *Comamonadaceae* as bacterial families describing arterial plaque. Saliva was also characterised on component 2 by the same families of bacteria both reported by the Koren et al. (2011) and our microbiome signature in the HMP data set.

Our comparative analysis demonstrates that sPLS-DA not only produces reliable and consistent results across different sequencing platforms and datasets but is also able to identify key members of the microbial community.

## Discussion

Traditionally, unsupervised dimension reduction multivariate approaches for microbiome data such as PCoA use pairwise distances or dissimilarities calculated on count data to scale microbial community abundances.However, the output of such method is limited to the visualisation of patterns in the data only. Our framework did not propose such distances for various reasons. From a theoretical point of view and as discussed by Warton et al. (2012), distance-based analyses make implicit assumptions on the mean-variance relationship in count data that may not hold, with the consequence of possible misleading results. From a practical point of view, a multivariate projection based method applied on a *n* × *n* similarity matrix does not allow us to identify bacteria driving differences between habitats. We therefore proposed to directly handle abundance data to achieve that goal.

In our study, we acknowledged that there was no clear consensus on the normalisation technique to apply for 16S OTU count data. The TSS normalisation is a popular approach to accommodate for varying sampling and sequencing depth White et al. (2009); Segata et al. (2011); Costello et al. (2015) but TSS produces compositional data which may lead to spurious results when applying traditional statistical methods Fernandes et al. (2014); Kurtz et al. (2015). Transforming compositional data using log ratios beforehand allows to circumvent this issue with either Isometric Log Ratio (ILR) or Centered Log Ratio transformation (CLR) Aitchison (1982); Filzmoser et al. (2009). We included those transformations as part of our framework to visualise diversity patterns (PCA) or to perform discriminant analysis and identify indicator species explaining abundance differences between samples (sPLS-DA). The multivariate projection-based methods that we propose to apply have the unique advantage of producing insightful graphical outputs for data interpretation but require appropriate log ratio transformation when dealing with compositional data. We applied the ILR transformation for PCA, as proposed by Filzmoser et al. (2009); Kalivodová et al. (2015) to overcome the CLR limitation that may lead to singular covariance matrices. For sPLS-DA however, the feature selection process requires *n* × *p* input matrix in order to identify indicator species. The advantage of using PLS methods and variants is their ability to reduce the dimension of the data, leading to non-singular matrix Boulesteix and Strimmer (2007). We therefore resorted to a CLR one-to-one transformation and showed that sPLS-DA delivered relevant results in our two case studies using TSS-CLR transformed data.

Our study also assessed the impact of normalisation using either TSS or the CSS normalisation recently proposed by Paulson et al. (2013) as an attractive method to account for sparse counts. In the Most Diverse case study we showed that both normalisations identified the same bacteria families. In the more complex Oral case study we observed differences as TSS-CLR identified more families than CSS. The data analyst must therefore keep in mind that normalisation is data specific and needs to be carefully chosen prior to statistical analysis.

Our framework proposes to handle repeated-measures design with a multilevel variance decomposition. This additional transformation step can also be seen as a scaling transformation to extract subtle differences between body sites or habitats within the same individuals. We anticipate that such experimental designs will become widely adopted in microbiome studies. Note however that our framework can also be used in a more general case with non repeated measures experiments.

As shown in Figure 1, **`mixMC`** proposes more extensive analytical features than univariate methods. When we compared the univariate and multivariate methods, we we found that the overall structure of the signatures were similar at the family level. However, dimension reduction multivariate approaches provide intuitive plots and numerical outputs for a better understanding of the discriminative ability of the OTU features identified.

Recently, a number of studies have investigated the link between gut and oral microbial communities Franzosa et al. (2014); Koren et al. (2011). In particular, Franzosa et al. (2014) showed that a subset of abundant oral microbes that are surviving transit to the gut are being linked with disease markers of atherosclerosis such as cholesterol Koren et al. (2011). From our detailed analyses, we reached similar conclusions identifying bacteria such as *Fusobacterium*, *Propionibacterium*, *Veillonella* in both the oral body sites from both HMP data sets (including plaque, tongue and gingiva) and stool microbiomes as underlined by Koren et al. (2011). Our comparative study with the Koren data set demonstrated that our multivariate method was able to identify a microbiome signature consistent across different individual cohorts and sequencing platforms. In addition, the results obtained by Koren et al. (2011) and our microbiome signatures identified from the most diverse HMP data set highlighted that microbial communities can not be considered discrete environments, but are, in fact, fluid environments.

## Conclusions

**`mixMC`** is a statistical analysis framework enabling holistic understanding of microbial communities. In this paper, we demonstrated the advantages of using multivariate methodologies for the statistical analysis of 16S compositional data, to summarise and reduce the dimension of possibly large data sets; to obtain a better understanding of the microbial communities through insightful graphical outputs; and to highlight features characterising and discriminating different environments. While our study has particularly focused on repeated-measures designs, the multivariate approach that we propose is not restricted to such designs only. Similar analyses can be performed on non-repeated designs to highlight relevant microbial features.

The multivariate approach sPLS-DA is a specific case of a larger family of projection-based multi-variate approaches, some of which also allow integration of different types of data. Our proposed analysis framework therefore paves the transition towards a ‘microbiome system biology’ approach by integrating large scale multi-‘omics studies such as metatranscriptomics, metabolomics or metaproteomics currently being collected by the integrative HMP project (Human Microbiome Project Consortium, 2014), therefore enabling the improvement of our understanding of the biomolecular activities and regulatory systems of human microbiota.

## Availability of supporting data

The data sets supporting the results of this article are available from the NIH Human Microbiome Project http://hmpdacc.org/HMQCP/all/ in raw data format, and in processed format on our website www.mixOmics.org/mixMC. R scripts and functions, together with a full tutorial and report to reproduce the results from the proposed framework are also available on our website.

## List of abbreviations

16S rRNA: - 16S ribosomal RNA
CLR: - Centered Log Ratio
CSS: - Cumulative Sum Scaling
HMP: - Human Microbiome Project
ILR: - Isometric Log Ratio
OTU: - Operational Taxonomy Unit
PCA: - Principal Component Analysis
PCoA: - Principal Coordinate Analysis
sPLS-DA: - sparse Partial Least Squares Discriminant Analysis
TSS: - Total Sum Scaling
ZIG: - Zero Inflated Gaussian

## Author′s contributions

KALC developed the methodologies, implemented the approaches, performed the statistical analyses and wrote the manuscript. MEC analysed the results and wrote the manuscript. XYC, FB and VAL helped with the implementation of the graphical outputs. FB and VL assisted with the R implementation of the framework and the tutorials on our website. RB ran the statistical analyses and participated in the design of the study. PR participated in the design of the study.

## Acknowledgements

KALC was supported in part by the Australian Cancer Research Foundation (ACRF) for the Diamantina Individualised Oncology Care Centre at The University of Queensland Diamantina Institute and the National Health and Medical Research Council (NHMRC) Career Development fellowship (APP1087415).The authors would like to thank Christian Cherveaux (Danone Nutricia Research) for fruitful discussions in the early stages of the project.

## Supplementary Material

**Table S1:** Description of the two HMP data sets through the preprocessing and normalization steps.

|  | Most Diverse body sites | Oral body sites |
| --- | --- | --- |
| # initial OTU | 43,140 | 43,140 |
| # raw OTU after pre-filtering | 1,674 | 1,562 |
| # samples | 162 | 657 |
| # unique individuals | 54 | 73 |
| # body sites | 3 | 9 |

## Electronic Supporting Information Legends

### Diverse, Oral and Koren TSS-CLR data: selected OTU

Contribution of selected OTU for each sPLS-DA component available electronically.

**Table S2:**
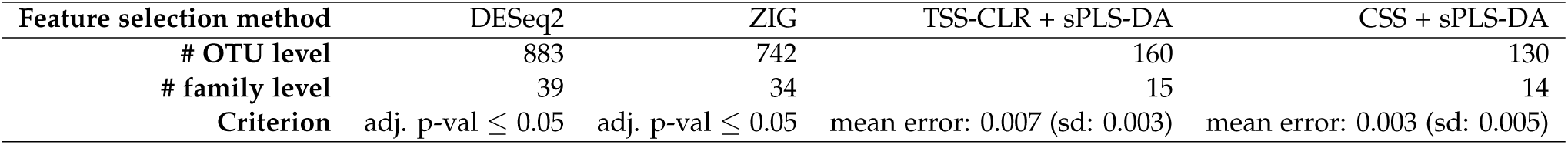
**Most diverse data, number of features selected by the different univariate and multivariate approaches at the OTU or family level**. The OTU features selection is based on either 5% significance level (adjusted FDR p-values) for DESeq2 and ZIG or the best classification performance with mean error rate across 10-fold cross-validation repeated 100 times (standard deviation) for sPLS-DA with two components.

**Table S3:** Oral data, performance of sPLS-DA per component and body site (TSS-CLR data). The mean classification error rate across 10-fold cross validation performed 100 times is indicated.

|  | comp 1 | comp 2 | comp 3 | comp 4 | comp 5 | comp 6 | comp 7 | comp 8 |
| --- | --- | --- | --- | --- | --- | --- | --- | --- |
| Attached Keratinized Gingiva | 1.00000 | 0.02603 | 0.09041 | 0.10164 | 0.10630 | 0.10795 | 0.11205 | 0.08411 |
| Buccal Mucosa | 1.00000 | 1.00000 | 0.19151 | 0.19973 | 0.21753 | 0.24685 | 0.23397 | 0.27151 |
| Hard Palate | 1.00000 | 0.24795 | 0.21397 | 0.32767 | 0.34740 | 0.10877 | 0.15370 | 0.14301 |
| Palatine Tonsils | 1.00000 | 1.00000 | 1.00000 | 1.00000 | 0.70959 | 0.70356 | 0.73890 | 0.53315 |
| Saliva | 1.00000 | 1.00000 | 1.00000 | 0.04493 | 0.04712 | 0.06137 | 0.06877 | 0.08603 |
| Subgingival Plaque | 1.00000 | 1.00000 | 0.99836 | 0.99123 | 0.38740 | 0.38466 | 0.39260 | 0.39288 |
| Supragingival Plaque | 0.00000 | 0.01370 | 0.01507 | 0.01945 | 0.20301 | 0.20384 | 0.19836 | 0.19315 |
| Throat | 1.00000 | 0.97425 | 0.99562 | 0.99644 | 0.99973 | 0.99808 | 0.53671 | 0.54630 |
| Tongue Dorsum | 0.00000 | 0.00055 | 0.00712 | 0.00000 | 0.01397 | 0.01425 | 0.09205 | 0.08904 |

**Table S4:**
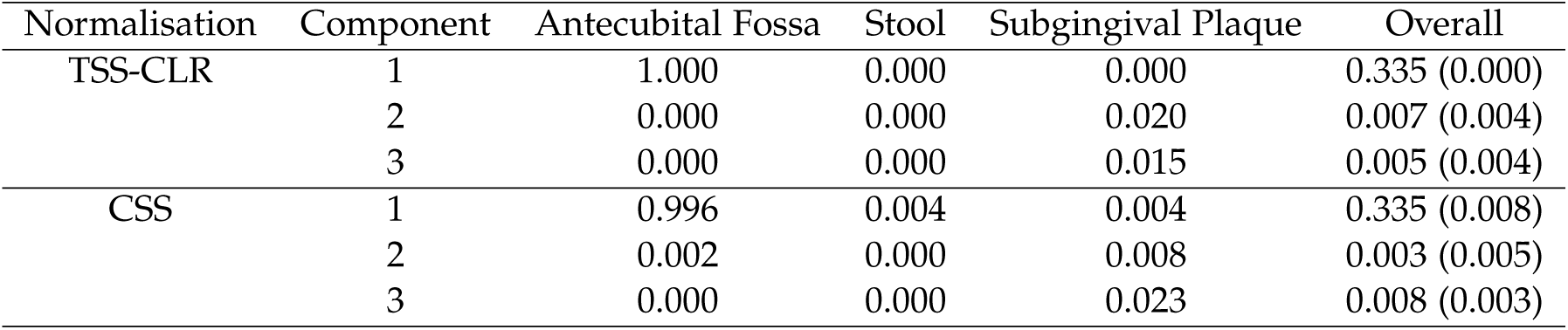
Most diverse data, performance of sPLS-DA per body site. Componentwise 100* 10-fold crossvalidation classification error rate for sPLS-DA applied to either TSS-CLR or CSS normalised counts with respect to each body site class leading to the optimal microbiome signature.

**Table S5:** Most diverse data, number of features contributing to each body site for each sPLS-DA component. The sPLS-DA model was applied to either TSS-CLR or CSS normalised counts. Contribution is defined as the body site for which the maximum median normalised OTU abundance is achieved at the OTU (family) level.

| Normalisation | Component | Antecubital Fossa | Stool | Subgingival Plaque | Total |
| --- | --- | --- | --- | --- | --- |
| TSS-CLR | 1 | 0 | 0 | 10 (5) | 10 (5) |
|  | 2 | 121 (5) | 29 (5) | 0 | 150 (10) |
| CSS | 1 | 0 | 0 | 10 (6) | 10 (6) |
|  | 2 | 60 (3) | 60 (5) | 0 | 120 (8) |

**Figure S1:**
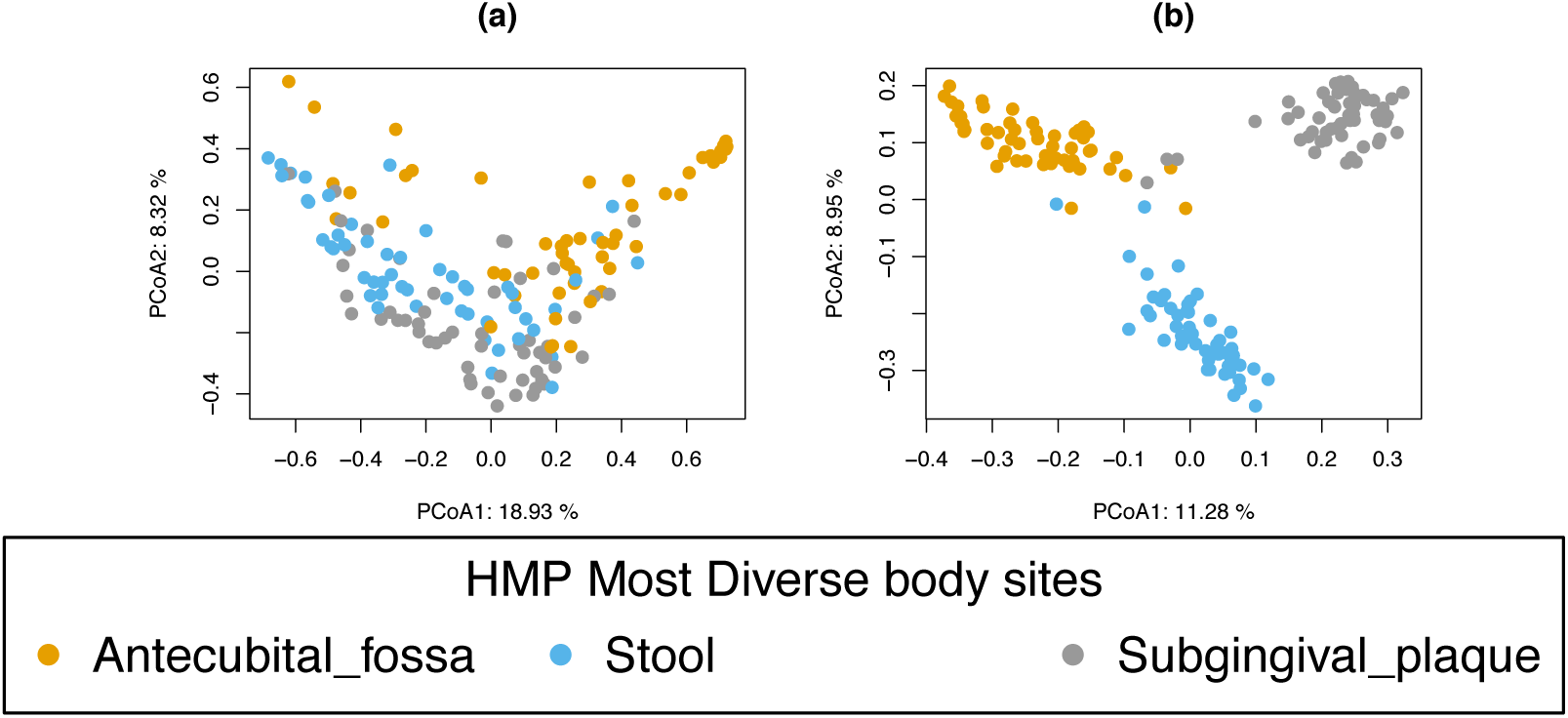
Most diverse data, PCoA sample plots. Sample plot on the first two coordinates with (**a**) weighted Unifrac (**b**) unweighted Unifrac calculated on the unfiltered OTU count table (based on 43,146 OTU).

**Figure S2:**
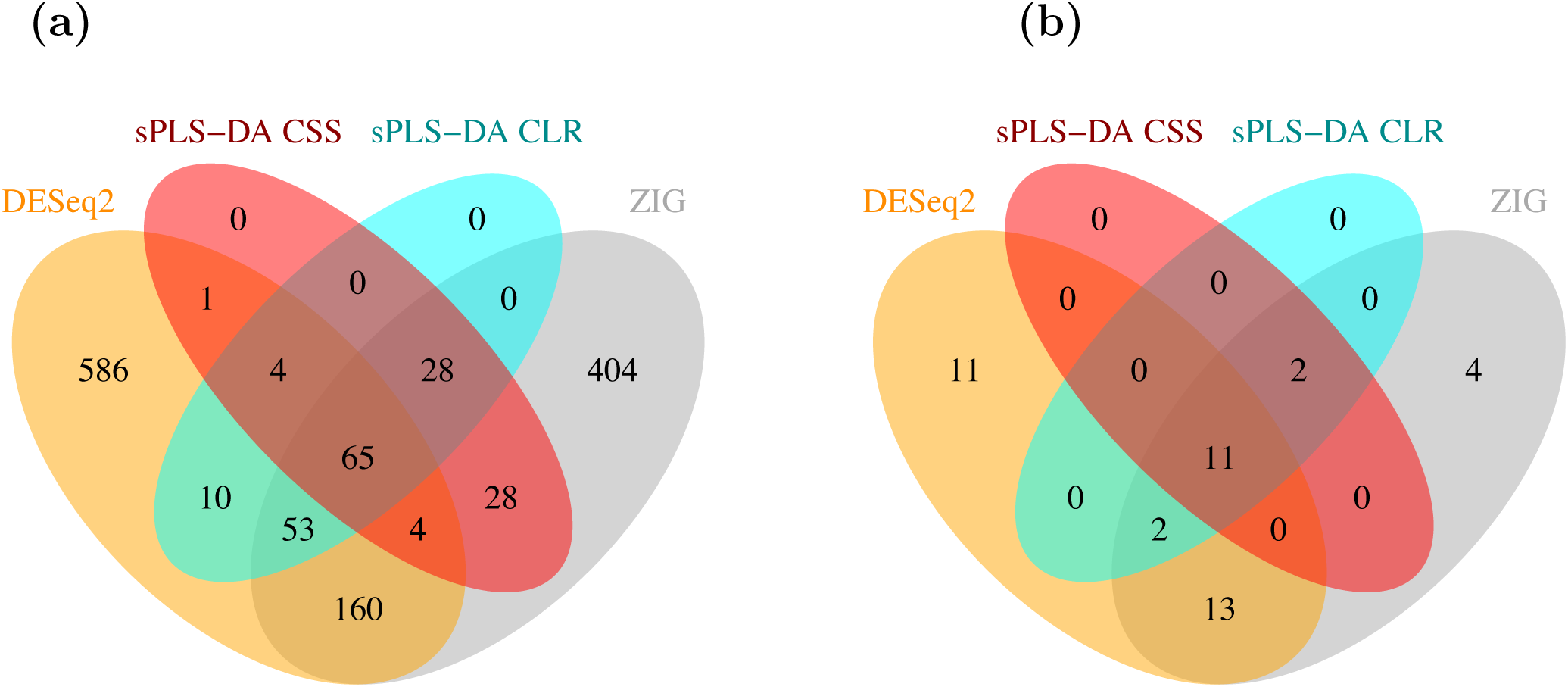
Most diverse data, comparison between univariate OTU selections and multivariate sPLS-DA selection. Comparison of the most differentially abundant features identified by DESeq2 and ZIG (FDR ≤ 0.05) and the most discriminative features identified by TSS-CLR+sPLS-DA or CSS+sPLS-DA (lowest mean classification error rate achieved when performing 100 * 10–fold cross-validation). (**a**): selection size at OTU level, (**b**): at the family level.

**Figure S3:**
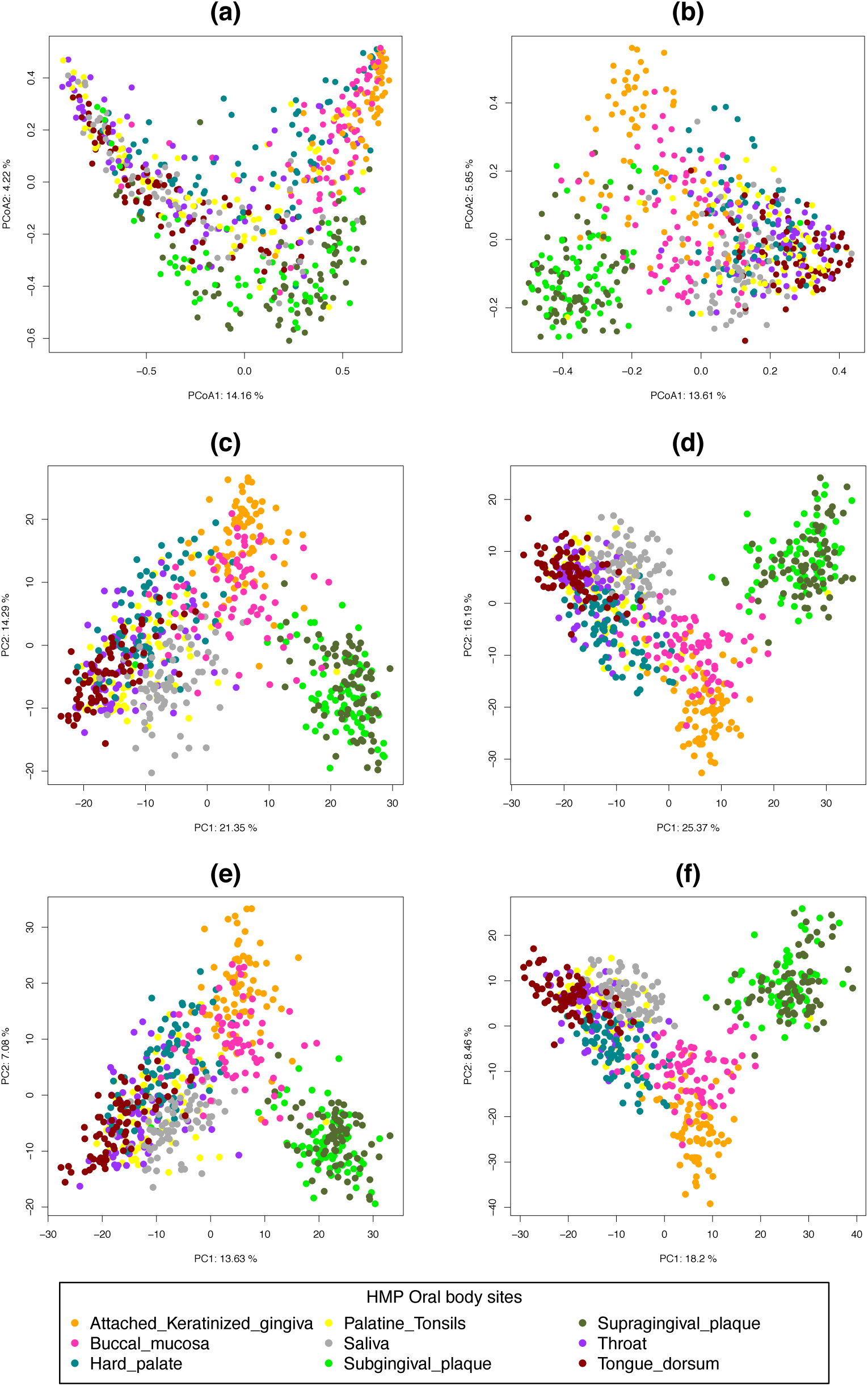
Oral data, PCoA and PCA sample plots. Sample plot on the first two coordinates with (**a**) weighted Unifrac (**b**) unweighted Unifrac calculated on the filtered OTU count table and on the first components 26 for (**c**) TSS-ILR and (**d**) TSS-ILR multilevel normalised OTU counts, and (**e**) CSS and (**f**) CSS multilevel normalised OTU counts. Colours indicate body sites and the percentage of explained variance for each principal component is indicated in the axes labels.

**Figure S4:**
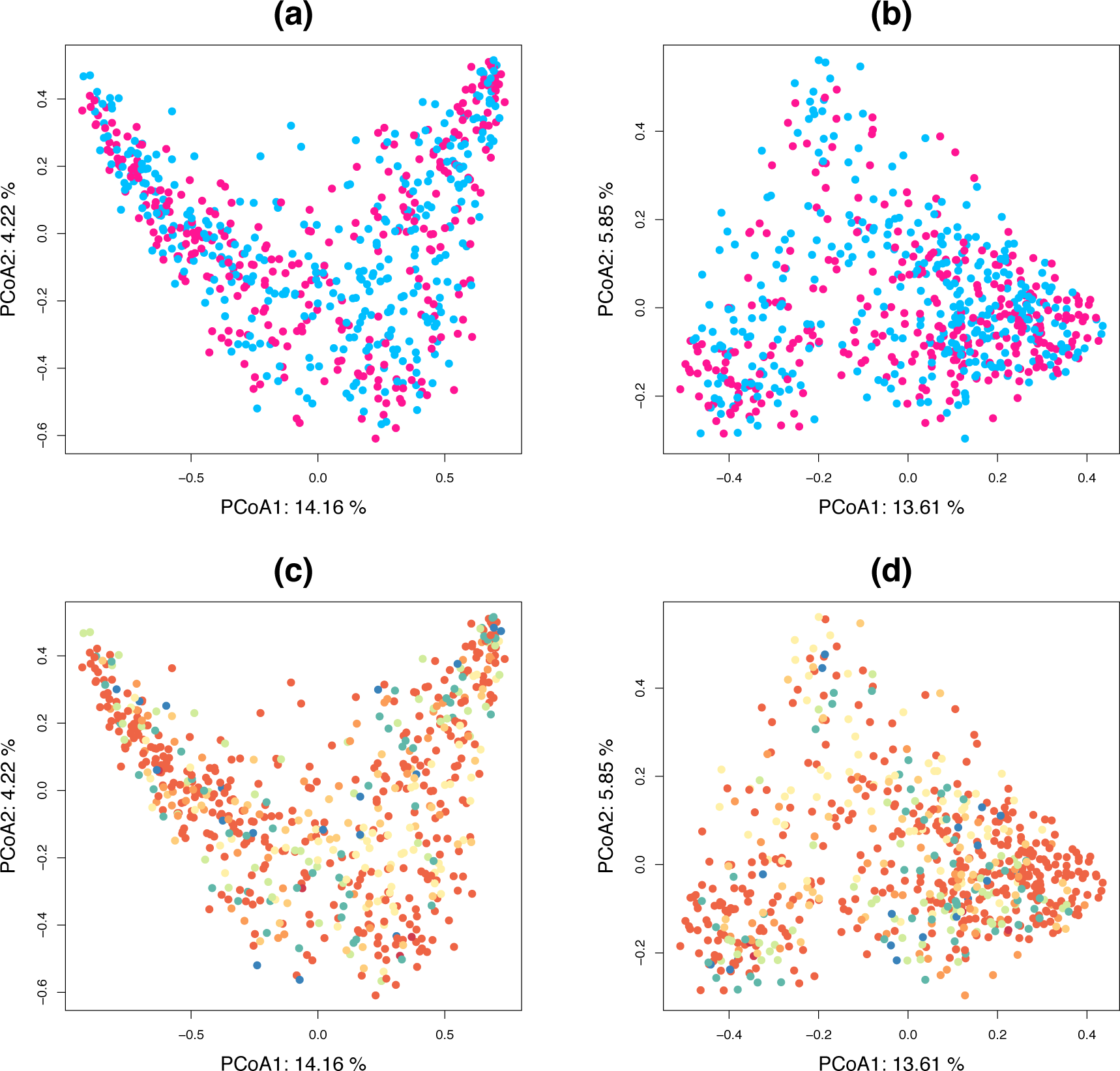
Oral data, PCoA sample plots with colors indicating gender or run centers. Sample plot on the first two coordinates with colors indicating gender in (**a**) weighted Unifrac or (**b**) unweighted Unifrac, or colors indicating run centers in(**c**) weighted Unifrac or (**d**) unweighted Unifrac calculated on the filtered OTU count table.

**Figure S5:**
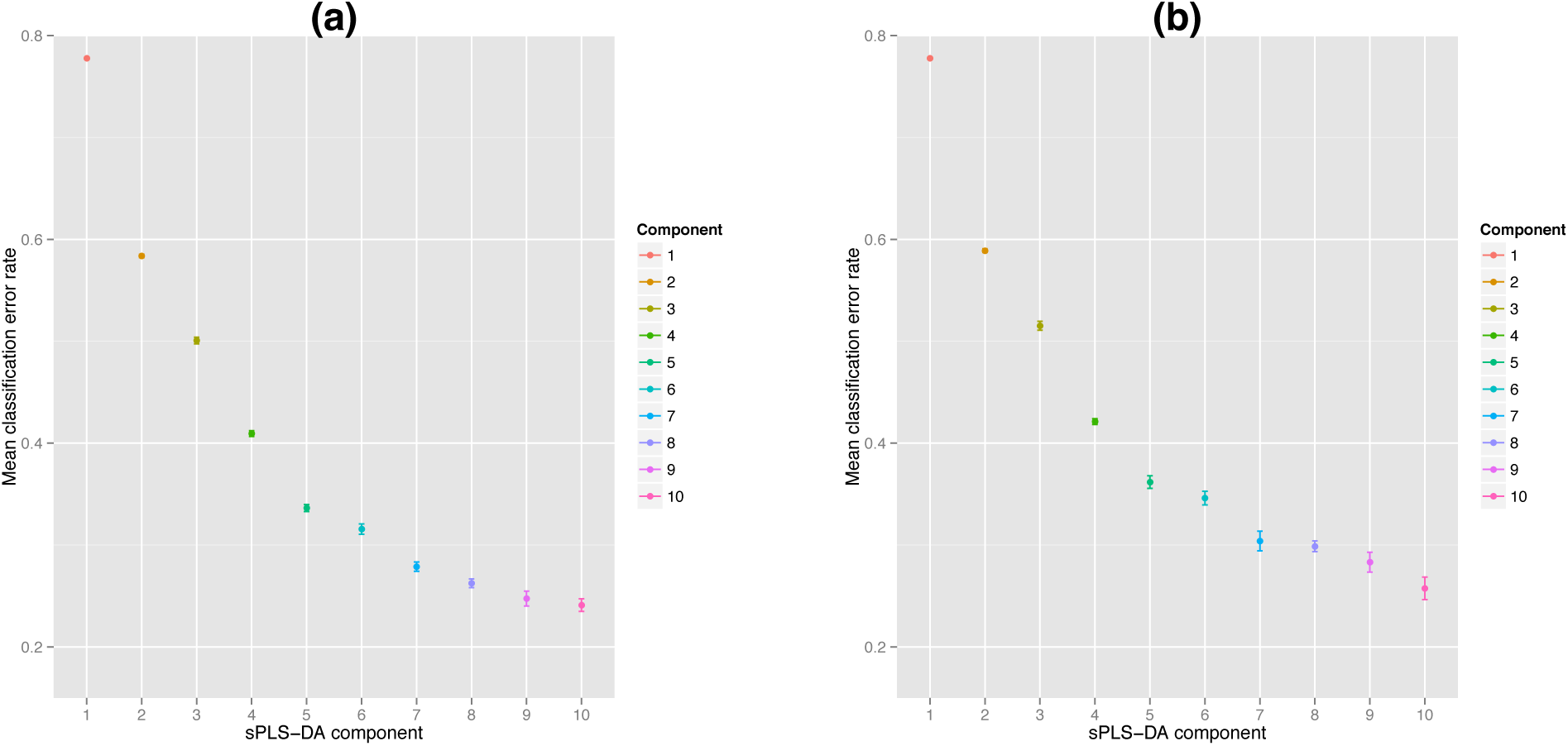
Oral data, sPLS-DA performance. Mean classification performance using 100 * 10 ‐fold cross-validation.Each component is based on an optimal selection of OTU features that leads to the best classification performance. The sPLS-DA classifier was applied on (**a**) TSS-CLR or (**b**) CSS normalised data.

**Figure S6:**
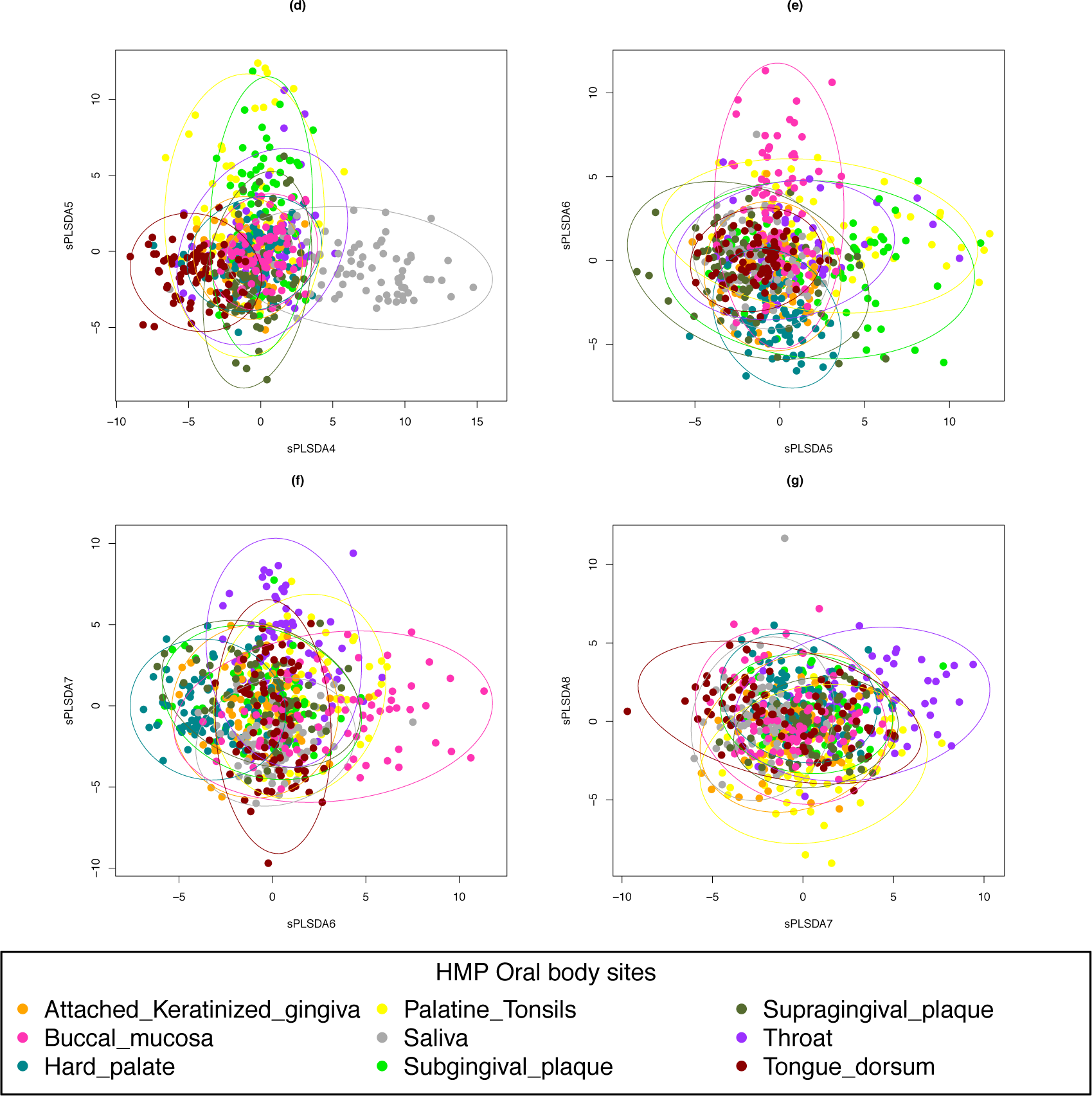
Oral data, sPLS-DA sample representation for the different components of the model. (**d**) Component 4 vs Component 5, (**e**) Component 5 vs Component 6, (**f**) Component 6 vs Component 7, (**g**) Component 7 vs Component 8. Colours indicate the different body sites, 95% confidence ellipses are displayed.

**Figure S7:**
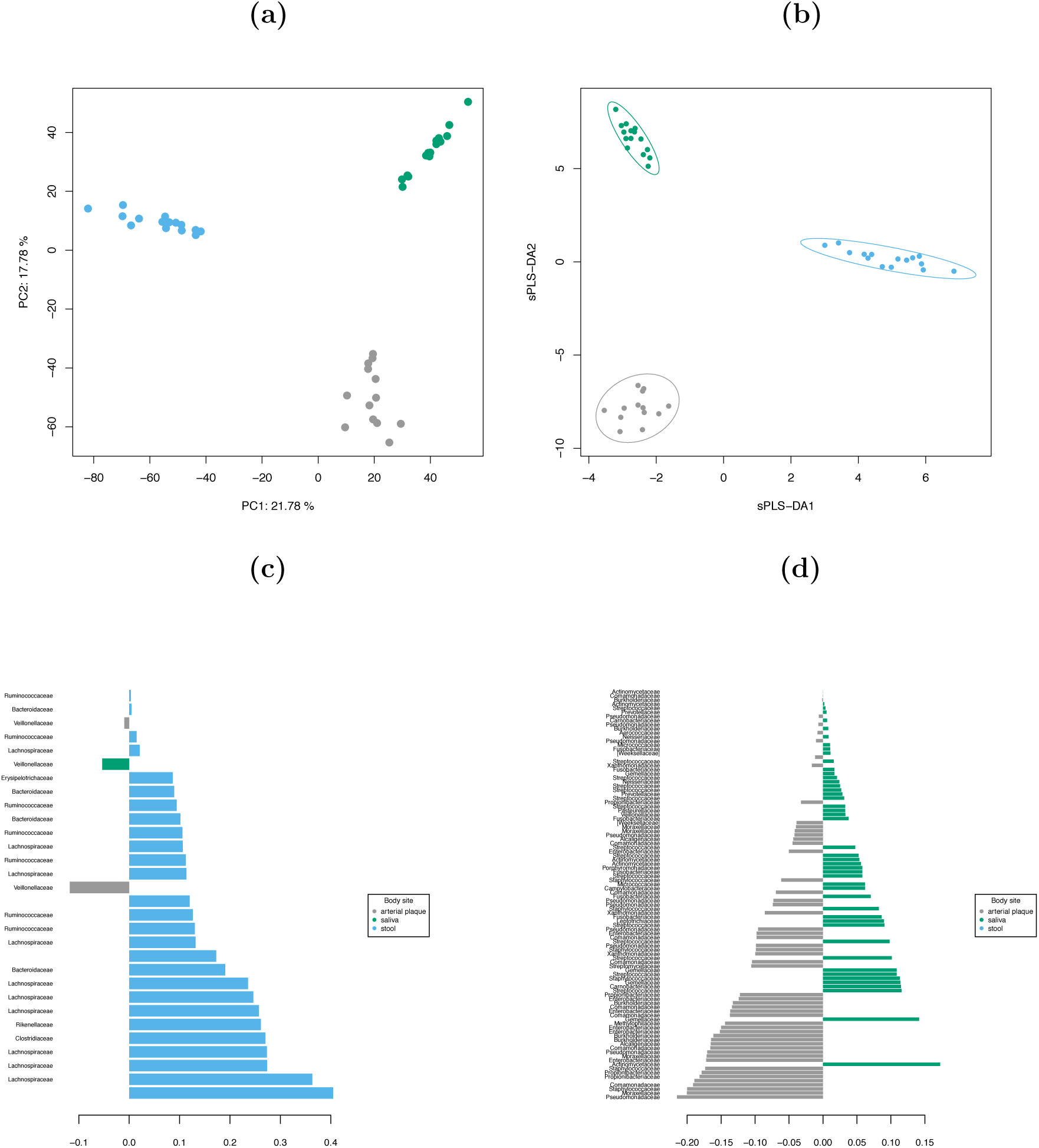
Koren data. Sample plot on the first two components with (**a**) PCA-ILR (**b**) sPLS-DA on selected OTU. Contribution plots on the (**c**) first component (30 OTU selected) and on the (**d**) second component (100 OTU selected). Colors indicate the body sites.

## References

Jill E Clarridge. Impact of 16S rRNA gene sequence analysis for identification of bacteria on clinical microbiology and infectious diseases. Clinical microbiology reviews, 17(4):840–862, 2004.

Susan M Huse, David Mark Welch, Hilary G Morrison, and Mitchell L Sogin. Ironing out the wrinkles in the rare biosphere through improved otu clustering. Environmental microbiology, 12 (7):1889–1898, 2010.

James Robert White, Niranjan Nagarajan, and Mihai Pop. Statistical methods for detecting differentially abundant features in clinical metagenomic samples. PLoS computational biology, 5 (4):e1000352, 2009.

Peter J Turnbaugh, Fredrik Bäckhed, Lucinda Fulton, and Jeffrey I Gordon. Diet-induced obesity is linked to marked but reversible alterations in the mouse distal gut microbiome. Cell host & microbe, 3(4):213–223, 2008.

Peter J Turnbaugh, Micah Hamady, Tanya Yatsunenko, Brandi L Cantarel, Alexis Duncan, Ruth E Ley, Mitchell L Sogin, William J Jones, Bruce A Roe, Jason P Affourtit, et al. A core gut microbiome in obese and lean twins. nature, 457(7228):480–484, 2009.

Sylvia H Duncan, GE Lobley, G Holtrop, J Ince, AM Johnstone, P Louis, and HJ Flint. Human colonic microbiota associated with diet, obesity and weight loss. International journal of obesity, 32(11):1720–1724, 2008.

Dirk Gevers, Subra Kugathasan, Lee A Denson, Yoshiki Vázquez-Baeza, Will Van Treuren, Boyu Ren, Emma Schwager, Dan Knights, Se Jin Song, Moran Yassour, et al. The treatment-naive microbiome in new-onset crohnâAZs disease. Cell host & microbe, 15(3):382–392, 2014.

Mary-Ellen Costello, Francesco Ciccia, Dana Willner, Nicole Warrington, Philip C Robinson, Brooke Gardiner, Mhairi Marshall, Tony J Kenna, Giovanni Triolo, and Matthew A Brown. Intestinal dysbiosis in ankylosing spondylitis. Arthritis & Rheumatology, 67(3):686âĂŞ691, 2015.

Mark D Robinson and Gordon K Smyth. Small-sample estimation of negative binomial dispersion, with applications to sage data. Biostatistics, 9(2):321–332, 2008.

Simon Anders and Wolfgang Huber. Differential expression analysis for sequence count data. Genome biol, 11(10):R106, 2010.

Michael I Love, Wolfgang Huber, and Simon Anders. Moderated estimation of fold change and dispersion for rna-seq data with deseq2. Genome biology, 15(12):550, 2014.

Paul J McMurdie and Susan Holmes. Waste not, want not: why rarefying microbiome data is inadmissible. PLoS computational biology, 10(4):e1003531, 2014.

Joseph N Paulson, O Colin Stine, Héctor Corrada Bravo, and Mihai Pop. Differential abundance analysis for microbial marker-gene surveys. Nature methods, 10(12):1200–1202, 2013.

John Aitchison. The statistical analysis of compositional data. Journal of the Royal Statistical Society. Series B (Methodological), pages 139–177,1982.

David Lovell, Vera Pawlowsky-Glahn, Juan José Egozcue, Samuel Marguerat, and Jürg Bähler. Proportionality: a valid alternative to correlation for relative data. PLoS computational biology, 11 (3):e1004075, 2015.

Yuguang Ban, Lingling An, and Hongmei Jiang. Investigating microbial co-occurrence patterns based on metagenomic compositional data. Bioinformatics, 31(20):3322–3329, 2015.

Siddhartha Mandal, Will Van Treuren, Richard A White, Merete Eggesbø, Rob Knight, and Shyamal D Peddada. Analysis of composition of microbiomes: a novel method for studying microbial composition. Microbial Ecology in Health and Disease, 26, 2015.

Andrew D Fernandes, Jennifer NS Reid, Jean M Macklaim, Thomas A McMurrough, David R Edgell, and Gregory B Gloor. Unifying the analysis of high-throughput sequencing datasets: characterizing rna-seq, 16s rrna gene sequencing and selective growth experiments by compositional data analysis. Microbiome, 2(1):1–13, 2014.

Zachary D Kurtz, Christian L Mueller, Emily R Miraldi, Dan R Littman, Martin J Blaser, and Richard A Bonneau. Sparse and compositionally robust inference of microbial ecological networks. PLoS Comput Biol, 11(5), 2015. doi: 10.1371/journal.pcbi.1004226.

Alžběta Kalivodová, Karel Hron, Peter Filzmoser, Lukáš Najdekr, Hana Janečková, and Tomas Adam. Pls-da for compositional data with application to metabolomics. Journal of Chemometrics, 29(1):21–28, 2015.

Nicola Segata, Jacques Izard, Levi Waldron, Dirk Gevers, Larisa Miropolsky, Wendy S Garrett, and Curtis Huttenhower. Metagenomic biomarker discovery and explanation. Genome Biol, 12(6): R60, 2011.

J Roger Bray and John T Curtis. An ordination of the upland forest communities of southern wisconsin. Ecological monographs, 27(4):325–349, 1957.

Catherine Lozupone and Rob Knight. Unifrac: a new phylogenetic method for comparing microbial communities. Applied and environmental microbiology, 71(12):8228–8235, 2005.

Catherine A Lozupone, Micah Hamady, Scott T Kelley, and Rob Knight. Quantitative and qualitative β diversity measures lead to different insights into factors that structure microbial communities. Applied and environmental microbiology, 73(5):1576–1585, 2007.

John C Gower. Principal coordinates analysis. Wiley StatsRef: Statistics Reference Online, 1998.

S Dolédec and D Chessel. Rythmes saisonniers et composantes stationnelles en milieu aquatique. i: Description d’un plan d’observation complet par projection de variables. Acta oecologica. Oecologia generalis, 8(3):403–426,1987.

Human Microbiome Project Consortium. A framework for human microbiome research. Nature, 486(7402):215–221, 2012a.

Human Microbiome Project Consortium. Structure, function and diversity of the healthy human microbiome. Nature, 486(7402):207–214, 2012b.

Omry Koren, Aymé Spor, Jenny Felin, Frida Fåk, Jesse Stombaugh, Valentina Tremaroli, Carl Johan Behre, Rob Knight, Björn Fagerberg, Ruth E Ley, et al. Human oral, gut, and plaque microbiota in patients with atherosclerosis. Proceedings of the National Academy of Sciences, 108(Supplement 1):4592–4598, 2011.

J Gregory Caporaso, Justin Kuczynski, Jesse Stombaugh, Kyle Bittinger, Frederic D Bushman, Elizabeth K Costello, Noah Fierer, Antonio Gonzalez Pena, Julia K Goodrich, Jeffrey I Gordon, et al. Qiime allows analysis of high-throughput community sequencing data. Nature methods, 7 (5):335–336, 2010.

Nicholas A Bokulich, Sathish Subramanian, Jeremiah J Faith, Dirk Gevers, Jeffrey I Gordon, Rob Knight, David A Mills, and J Gregory Caporaso. Quality-filtering vastly improves diversity estimates from illumina amplicon sequencing. Nature methods, 10(1):57–59, 2013.

Dan Knights, Laura Wegener Parfrey, Jesse Zaneveld, Catherine Lozupone, and Rob Knight. Human-associated microbial signatures: examining their predictive value. Cell host & microbe, 10 (4):292–296, 2011.

Manimozhiyan Arumugam, Jeroen Raes, Eric Pelletier, Denis Le Paslier, Takuji Yamada, Daniel R Mende, Gabriel R Fernandes, Julien Tap, Thomas Bruls, Jean-Michel Batto, et al. Enterotypes of the human gut microbiome. nature, 473(7346):174–180, 2011.

Joseph Nathaniel Paulson, M Pop, and Hector Corrada Bravo. metagenomeseq: Statistical analysis for sparse high-throughput sequencing, 2015. URL http://cbcb.umd.edu/software/metagenomeSeq Bioconductor package: 1.6.0.

Peter Filzmoser, Karel Hron, and Clemens Reimann. Principal component analysis for compositional data with outliers. Environmetrics, 20(6):621–632, 2009.

Juan José Egozcue, Vera Pawlowsky-Glahn, Glòria Mateu-Figueras, and Carles Barcelo-Vidal. Isometric logratio transformations for compositional data analysis. Mathematical Geology, 35(3): 279–300, 2003.

Matthias Templ, Karel Hron, and Peter Filzmoser. robCompositions: an r-package for robust statistical analysis of compositional data, 2011.

Johan A Westerhuis, Ewoud JJ van Velzen, Huub CJ Hoefsloot, and Age K Smilde. Multivariate paired data analysis: multilevel plsda versus oplsda. Metabolomics, 6(1):119–128, 2010.

Benoit Liquet, Kim-Anh Lê Cao, Hakim Hocini, and Rodolphe Thiébaut. A novel approach for biomarker selection and the integration of repeated measures experiments from two assays. BMC bioinformatics, 13(1):325, 2012.

Jasmin Straube, Alain-Dominique Gorse, Bevan Emma Huang, Kim-Anh Lê Cao, et al. A linear mixed model spline framework for analysing time course ?omics? data. PloS one, 10(8):e0134540, 2015.

Kim-Anh Lê Cao, Simon Boitard, and Philippe Besse. Sparse PLS Discriminant Analysis: biologically relevant feature selection and graphical displays for multiclass problems. BMC bioinformatics, 12(1):253, 2011.

Svante Wold, Michael Sjöström, and Lennart Eriksson. Pls-regression: a basic tool of chemometrics. Chemometrics and intelligent laboratory systems, 58(2):109–130, 2001.

Robert Tibshirani. Regression shrinkage and selection via the lasso. Journal of the Royal Statistical Society. Series B (Methodological), pages 267–288, 1996.

F Asnicar, G Weingart, TL Tickle, C Huttenhower, and N Segata. Compact graphical representation of phylogenetic data and metadata with graphlan. PeerJ, 2015. doi: https://dx.doi.org/10.7717/peerj.1029.

K.-A Lê Cao, I. González, S. Déjean, F. Rohart, and B. Gautier. mixomics: Omics data integration project.

Yoav Benjamini and Yosef Hochberg. Controlling the false discovery rate: a practical and powerful approach to multiple testing. Journal of the Royal Statistical Society. Series B (Methodological), pages 289–300, 1995.

Kelvin Li, Monika Bihan, Shibu Yooseph, and Barbara A Methé. Analyses of the microbial diversity across the human microbiome. PloS one, 7(6):e32118, 2012.

Human Microbiome Project Consortium. Evaluation of 16S rDNA-based community profiling for human microbiome research. PLoS One, 7(6):e39315, 2012c.

Xuesong He, Jeffrey S McLean, Anna Edlund, Shibu Yooseph, Adam P Hall, Su-Yang Liu, Pieter C Dorrestein, Eduardo Esquenazi, Ryan C Hunter, Genhong Cheng, et al. Cultivation of a human-associated tm7 phylotype reveals a reduced genome and epibiotic parasitic lifestyle. Proceedings of the National Academy of Sciences, 112(1):244–249, 2015.

David I Warton, Stephen T Wright, and Yi Wang. Distance-based multivariate analyses confound location and dispersion effects. Methods in Ecology and Evolution, 3(1):89–101, 2012.

Anne-Laure Boulesteix and Korbinian Strimmer. Partial least squares: a versatile tool for the analysis of high-dimensional genomic data. Briefings in bioinformatics, 8(1):32–44, 2007.

Eric A Franzosa, Xochitl C Morgan, Nicola Segata, Levi Waldron, Joshua Reyes, Ashlee M Earl, Georgia Giannoukos, Matthew R Boylan, Dawn Ciulla, Dirk Gevers, et al. Relating the metatranscriptome and metagenome of the human gut. Proceedings of the National Academy of Sciences, 111(22):E2329–E2338, 2014.

Human Microbiome Project Consortium. The integrative human microbiome project: Dynamic analysis of microbiome-host omics profiles during periods of human health and disease. Cell host & microbe, 16(3):276, 2014.

